# ABCE1-dependent translational control links Fe-S cluster biogenesis to parasite growth and lipid homeostasis in *Toxoplasma gondii*

**DOI:** 10.64898/2026.07.01.735774

**Authors:** Ambre J.M. Maupin, Maxime Gonzalez Durany, Eléa A. Renaud, Arnault Graindorge, Vincent Demolombe, Laurence Berry, Valérie Rofidal, Sébastien Besteiro

## Abstract

*Toxoplasma gondii* relies on tightly regulated protein synthesis to adapt to diverse host environments and to progress through its developmental stages. Here, we investigated the role of the ATP-binding cassette protein ABCE1, a broadly-conserved factor involved in ribosome recycling and translational control. Using a conditional knockdown approach, we demonstrate that depletion of TgABCE1 severely impairs parasite growth and disrupts global protein synthesis, confirming its essential role in maintaining translational capacity. TgABCE1 function depends on the incorporation of iron-sulfur (Fe-S) clusters, likely mediated by the cytosolic iron-sulfur assembly (CIA) pathway component HCF101. Depletion of TgABCE1 phenocopies the defects observed in TgHCF101-depleted parasites, supporting a functional link between these proteins. Notably, loss of TgABCE1 also disrupts lipid homeostasis, resulting in the accumulation of lipid droplets. Together, these findings uncover a critical link between translational regulation, Fe-S cluster biogenesis, and lipid homeostasis, highlighting the central role of proteostasis in parasite survival and development.

## Introduction

*Toxoplasma gondii* is an obligate intracellular parasite of the phylum Apicomplexa, which also includes important human pathogens such as *Plasmodium* and *Cryptosporidium*. It can infect virtually all warm-blooded animals and has a complex lifecycle, with sexual reproduction occurring in felids (the definitive hosts) and asexual development taking place in intermediate hosts, including humans. In these hosts, the parasite transitions from rapidly replicating tachyzoites, responsible for acute infection, to slow-growing bradyzoites that form persistent tissue cysts, particularly in muscle and neural tissues. This capacity to establish long-term latent infection, with the potential for reactivation under immunosuppression, is central to its persistence and clinical importance^1^.

Given the various environments encountered by *T. gondii* during its lifecycle, protein synthesis and translational control are of particular importance, as they play a central role in regulating the parasite’s access to nutrients and host-derived resources. These processes are not only critical for adaptation to environmental and cellular stresses^2,3^, but also for orchestrating the expression of stage-specific proteomes underlying its developmental transitions^4,5^.

One central and highly conserved regulator of protein synthesis is the ATP-binding cassette (ABC) ATPase ABCE1 (also known as Rli1 in yeast). It functions at the interface of translation initiation, termination, and ribosome recycling^6^. By promoting the dissociation of post-termination ribosomal complexes into their subunits, ABCE1 ensures efficient ribosome recycling and maintains the pool of active ribosomes available for new rounds of translation^7,8^. ABCE1 was also found associated with eukaryotic initiation factors and as a part of ribosomal pre-initiation complexes^9–11^. Studies across several eukaryotes have shown that ABCE1 depletion disrupts multiple stages of the translation cycle and perturbs cellular protein homeostasis^12,13^. Through this role, this important protein acts as a key factor sustaining translational capacity and fidelity, and thereby supports rapid proteome remodeling in response to cellular demands.

Unlike most ABC proteins, members of the ABCE subfamily lack membrane-spanning domains^14^. ABCE1 is further distinguished by the fact that it contains two iron-sulfur (Fe-S) clusters^15,16^. The Fe-S clusters of ABCE1 stabilize its N-terminal domain, which is essential for ribosome binding, and serve as a structural scaffold that conveys conformational changes from the ATPase domains to the ribosome-interaction surfaces^16,17^. This mechanism likely allows ABCE1 to couple ATP hydrolysis to the splitting of post-termination ribosomal complexes into subunits^18,19^.

ABCE1 acquires its Fe-S clusters via the cytosolic Fe-S cluster assembly (CIA) machinery, with the final transfer steps mediated by adaptor proteins such as Yae1-Lto1 in Opisthokonta^20^, including yeast and mammals. However, these adaptors are absent in Apicomplexa, where the protein HCF101 is instead likely responsible for delivering Fe-S clusters to ABCE1^21^. Many *T. gondii* Fe-S proteins are predicted to have housekeeping functions, with several potentially involved in regulating translation, including ABCE1 but also tRNA modifying proteins^22^. In fact, several steps of the translation process depend on iron-containing enzymes^23^ and, accordingly, global iron deprivation was recently shown to trigger translational repression in the parasite^24^.

Despite considerable advances over the past decade in identifying the key components involved in Fe-S cluster acquisition, the mechanisms by which the CIA pathway specifically recognizes its diverse target client Fe-S proteins remain unclear. Specific transfer proteins may mediate selective interactions with privileged client proteins^20^, and while the molecular determinants of these interactions are beginning to emerge^25,26^, they remain far from fully understood. ABCE1 was found to directly interact with HCF101 in *T. gondii*, and TgABCE1 levels were reduced in the TgHCF101 mutant, suggesting decreased stability due to defective Fe-S cluster incorporation^21^. However, whether TgABCE1 represents the primary client protein of TgHCF101 is still an open question.

To investigate this further, we generated a TgABCE1 conditional knockdown cell line. First, it allowed us to demonstrate for the first time that TgABCE1, as in other eukaryotes, is a key regulator of protein synthesis in *T. gondii*. Moreover, its depletion essentially phenocopies that of TgHCF101, potentially suggesting that TgABCE1 is a main client of TgHCF101. Finally, we established a link between perturbation of lipid homeostasis, manifested by the accumulation of lipid droplets (LDs, as previously observed under iron depletion^27^ and in CIA pathway mutants^21,28^), to TgABCE1 function disruption and protein synthesis inhibition. Our study provides novel insights into the importance of proteostasis for proper regulation of lipid metabolism and its implications for parasite growth.

## Results

### TgABCE1 is essential for the viability of *T. gondii* tachyzoites

To decipher TgABCE1 function, we generated a transgenic cell line in which we modified the 5′ region of the *TgABCE1* gene by homologous recombination to replace the endogenous promoter by an anhydrotetracycline- (ATc-) inducible Tet07SAG4 promoter, and at the same time adding a sequence coding for a N-terminal HA tag (**Fig. S1**). In this conditional knock-down (cKD) HA-TgABCE1 cell line, the addition of ATc can repress the transcription of *TgABCE1* thanks to a Tet-Off system^29^. Immunoblot analysis revealed an HA-tagged protein consistent with the predicted molecular mass of TgABCE1 (69 kDa), and showed that its depletion was efficient as early as 24 h after ATc addition (**Fig. 1A**). Immunofluorescence analysis (IFA) showed a cytoplasmic distribution of HA-tagged TgABCE1 and also confirmed the depletion of the protein upon ATc addition (**Fig. 1B**).

**Figure. 1.**
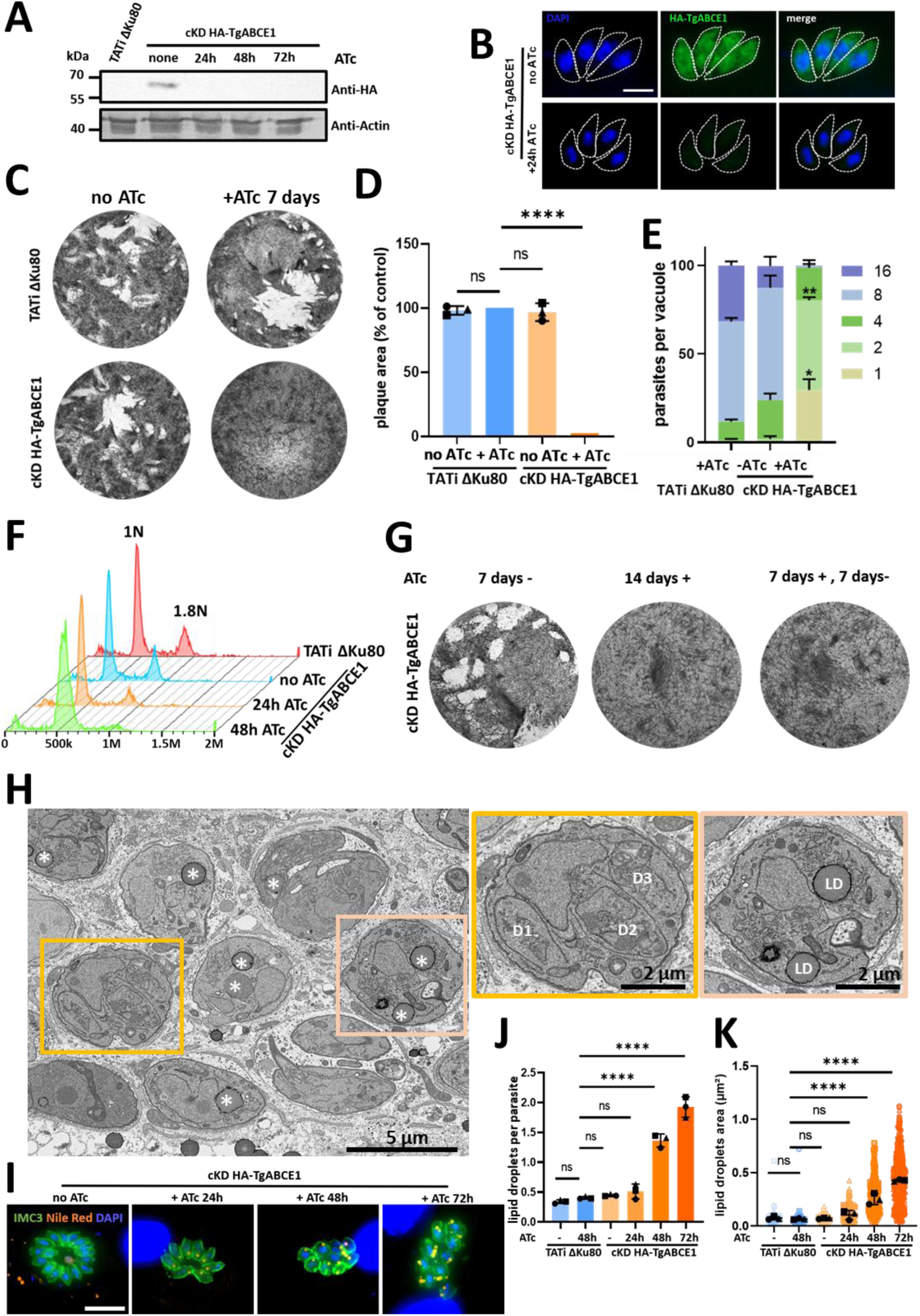
TgABCE1 is essential for the growth of *T. gondii*. **A.** Immunoblot analysis of the cKD HA-TgABCE1 mutant showing efficient tagging and down-regulation of TgABCE1 starting at 24 h of treatment with ATc. Actin was used as a loading control. **B.** Immunofluorescence assay showing cytosolic localization of the HA-tagged TgABCE1 protein, and total depletion of the protein after 24 h of ATc treatment. Parasites are outlined and DNA was stained with 4′,6-diamidino-2-phenylindole (DAPI). Scale bar = 5 μm. **C.** Plaque assays showing that TgABCE1 knockdown renders the parasite unable to form plaques in the host cell monolayer. These tests were carried out by infecting a monolayer of HFFs with the parental strain TATi ΔKu80 or cKD HA-TgABCE1 cell lines for 7 days in the presence or absence of ATc. **D.** Quantification of plaque area observed in (**C**). Results are expressed as the percentage of lysed area relative to the control (TATi ΔKu80 +ATc, set as 100% for reference). Values represented are mean ± SD from *n* = 3 independent biological replicates, ns: not significant (*p*-value >0.05), **** *p*-value ≤0.0001 from one-way ANOVA with Dunnett’s multiple comparison test. **E.** Replication assay of parental (TATi ΔKu80) and transgenic (cKD HA-TgABCE1) cell lines: parasites were pre-cultured for 24 h in the presence or absence of ATc and allowed to invade HFF-coated coverslips for another 24 h in the presence or absence of ATc, for a total of up to 48 h. The number of parasites per vacuole was quantified for each condition and expressed as a percentage; 100 vacuoles were counted for each condition. Values represented are mean ± SD of *n* = 3 independent biological replicates. Statistical significance was determined between each category by Student’s t-test and noted as follows: * *p*-value ≤0.05, ** *p*-value ≤0.01. F. DNA content analysis by flow cytometry on TATi ΔKu80 and cKD-TgABCE1 parasites treated or not with ATc up to 2 days and stained with propidium iodide. 1N and 1.8N represent the ploidy. **G.** Plaque assays were carried out with cKD HA-TgABCE1 parasites as described in (**C**), but for the “7 days+7 days–” condition, ATc was washed out after 7 days, and parasites were allowed to grow for another 7 days without ATc, while in the “14 days”’ condition, ATc treatment was maintained for 7 more days. The “7 days –” control was kept without ATc for 7 days of growth. **H.** Electron micrograph of cKD HA-TgABCE1 parasites pre-incubated with ATc for 24 h, then allowed to reinvade for an additional 24 h in the presence of ATc. Asterisks mark lipid droplets. The right panel shows a magnification of the selection on the left panel, highlighting defects in daughter cell budding within the mother cell, as well as the accumulation of lipid droplets. Pixel size: 5 nm. D: daughter bud, LD: lipid droplet. **I.** Fluorescent imaging of cKD HA-TgABCE1 parasites either untreated, or treated for 24 h, 48 h or 72 h with ATc, showing an accumulation of lipid droplets upon TgABCE1 depletion. LDs were detected with Nile Red (orange), parasites are outlined with an anti-IMC3 antibody (green), and DNA is stained with DAPI. Scale bar = 10 μm. **J, K.** Quantification of LD area and number, respectively. 100 parasites were analyzed per condition. The parental (TATi ΔKu80) and cKD HA-TgABCE1 transgenic parasites were grown in absence or presence of ATc for 24 h, 48 h, or 72 h. Values are represented as the mean ±standard deviation of *n* = 3 independent biological replicates (different symbols represent different series); ns: not significant (*p*-value >0.05), **** *p*-value ≤0.0001 using one-way ANOVA with Dunnett’s multiple comparison test. LD, lipid droplet.

We next evaluated the effect of TgABCE1 depletion on the lytic cycle of tachyzoites (the highly replicative and invasive stage responsible for acute infection) by assessing plaque formation on a monolayer of host cells. Host cells were infected and cultured with or without ATc for 7 days (**Fig. 1C, D**). TgABCE1 depletion completely abolished plaque formation. Next, cKD HA-TgABCE1 parasites were pretreated with ATc for 24 h, mechanically released, and used to infect fresh host cells for 24 h in ATc before parasite counting (**Fig. 1E**). ATc treatment led to vacuoles with reduced parasite numbers, suggesting that the lytic cycle defect reflected a replication problem. When evaluating DNA content through propidium iodide labeling and flow cytometer-based analysis, we observed that depletion of TgABCE1 lead to a progressive block in DNA replication, as indicated by a decrease in mitotic parasites (**Fig. 1F**). However, impairment of growth and replication do not necessarily indicate parasite death, as tachyzoites can differentiate into slow-growing, cyst-forming bradyzoites^30^. We then used *Dolichos biflorus* lectin (DBL), which specifically binds CST1^31^, a cyst wall glycoprotein synthesized early during stage conversion, and accumulating as differentiation progresses (**Fig. S2A**). We did not detect any sign of cystogenesis upon removal of TgABCE1. We next repeated the plaque assay with ATc removal after 7 days, followed by an additional 7-day culture before evaluating plaque formation (**Fig. 1G**). No plaque was observed after ATc removal, suggesting that the parasites were not able to recover after 7 days of TgABCE1 depletion. Together, these results indicate that TgABCE1 depletion causes parasite death, rather than simply slowing growth or promoting persistence.

We used electron microscopy to examine the effects of TgABCE1 depletion on developing parasites (**Fig. 1H**) and observed important defects in cytokinesis and daughter cell budding, similar to those seen upon TgHCF101 depletion^21^. Another striking feature shared with the TgHCF101 mutant was the accumulation of LDs. To quantify this, we used Nile Red, a fluorescent dye selective for neutral lipids, combined with microscopic imaging (**Fig. 1I, J, K**). We observed a marked increase in both the number and size of LDs in cKD HA-TgABCE1 parasites after up to 3 days of ATc treatment. We could observe that the induced LDs persisted in spite of ATc washout and further incubation for up to 48 h (**Fig. S2B, C, D**). In fact, immunoblot analysis showed that TgABCE1 expression was not restored following ATc washout (**Fig. S2E**), and that parasites remained largely unfit, as assessed by plaque assays, although still able to generate some small plaques (**Fig. S2F, G**).

Collectively, these data show that TgABCE1 depletion leads to an irreversible demise of the parasites, characterized by severe defects in parasite division and accompanied by the accumulation of LDs.

### Fe-S cluster importance for TgABCE1 function

ABCE1 typically consists of three main domains^32^: two nucleotide-binding domains (NBDs) arranged in a head-to-tail orientation via a flexible linker and hinge region, and present in all ABC proteins, as well as a N-terminal Fe-S binding domain (**Fig. 2A**, **Fig. S3**). The latter includes eight cysteine residues that coordinate two [4Fe-4S] clusters^16^. In order to assess the functional importance of Fe-S cluster coordination, we chose to complement the cKD HA-TgABCDE1 mutant with either a myc-tagged wild-type copy of the protein or a myc-tagged TgABCE1 variant mutated on cysteine 36 (C36S), which is part of the first Fe-S cluster coordination motif (**Fig. S4**). Both myc-tagged proteins displayed the expected cytoplasmic localization by IFA (**Fig. 2B**) and were constitutively expressed following depletion of the regulatable copy upon ATc addition, as confirmed by immunoblotting (**Fig. 2C**). Expression of the wild-type copy restored parasite fitness, as assessed by plaque assay (**Fig. 2D, E**), but in sharp contrast, the C36S mutant failed to rescue growth (**Fig. 2D**). Consistently, the LDs accumulation phenotype was reversed upon expression of wild-type TgABCE1, whereas the C36S mutant was unable to do so (**Fig. 2F, G, H**). Altogether, these results show that altering a single amino acid coordinating only one of the two Fe-S clusters of TgABCE1 is enough to abolish its function.

**Figure 2.**
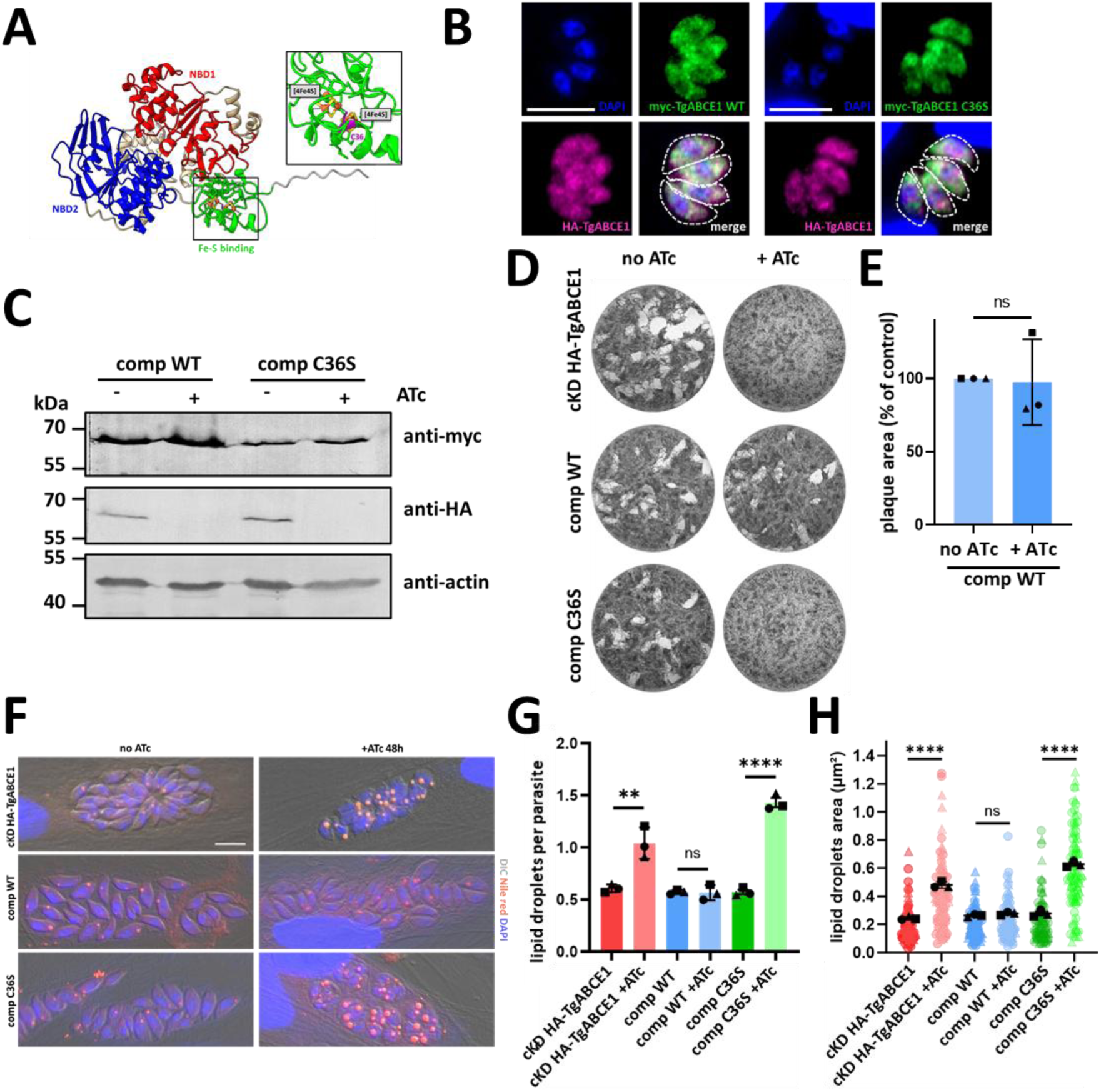
Key role of Fe-S in TgABCE1 function. **A.** AlphaFold-predicted structure of TgABCE1, highlighting the two nucleotide-binding domains (NBDs) (blue and red) and the Fe-S clusters binding region (green). The inset shows a zoom of the two Fe-S binding sites, highlighting cysteine 36 (pink) substituted with a serine in the comp C36S mutant. **B**. Immunofluorescence with anti-myc antibody confirms the cytoplasmic localization of the myc-tagged copies of TgABCE1. The parasite shapes are outlined. DNA was stained with DAPI. Scale bar = 5 μm. **C.** Immunoblot analysis showing the expression of the additional myc-tagged TgABCE1 copies and the depletion of the ATc-regulated HA-tagged copy TgABCE1 in the cKD HA-TgABCE1 comp WT/comp C36S cell lines. Actin is used as a loading control. **D.** Plaque assay showing restoration of growth by the additional WT copy upon depletion of the ATc-regulated copy, contrarily to the comp C36S mutant. Plaque assays were carried out by infecting a monolayer of HFFs with the cKD HA-TgABCE1, the cKD HA-TgABCE1 comp WT and the cKD HA-TgABCE1 comp C36S cell lines for 7 days in the presence or absence of ATc. **E.** Quantification of plaque area observed in (**D**). Results are expressed as a percentage of the lysed area relative to the control (cKD HA-TgABCE1 comp WT set as 100% for reference). Values represented are mean ± SD from *n* = 3 independent biological replicates, ns: not significant (>0.05) *p*-value from Student’s t-test. **F.** Representative images of vacuoles containing parasites grown continuously in the presence of ATc for 48 h and stained by Nile Red (orange) for lipid droplets. DIC: differential interference contrast. DNA was stained with DAPI. Scale bar = 5 µm. **G, H.** Quantification of LD number per parasite area, respectively; 100 parasites were analyzed per condition. The cKD HA-TgABCE1 comp WT or C36S parasites were grown in presence of ATc for 48 h. Values are represented as the mean ±standard deviation of *n* = 3 independent biological replicates (different symbols represent different series); ns, not significant (*p*-value >0.05), ** *p*-value ≤0.01, **** *p*-value ≤0.0001, Student’s t-test.

One peculiarity of ABCE1 from *T. gondii*, but also other apicomplexan parasites, compared with isoforms from plant or *Opisthokonta* is the presence of an N-terminal extension before the Fe-S cluster binding domain (**Fig. 3A, B and Fig. S3**). Interestingly, ABCE1 homologs of the *Apicomplexa*-relative *Chromera*, that also likely uses a cytosolic isoform of HCF101 for Fe-S cluster incorporation^21^, also possesses this lysine-rich N-terminal motif, which is absent from organisms having no HCF101 homolog, or at least no cytoplasmic isoform of the protein like land plants (**Fig. 3B, Fig. S3**). Therefore, we hypothesized that this sequence could be a major determinant of the TgHCF101/TgABCE1 interaction and could be key to Fe-S transfer. Thus, we complemented the cKD HA-TgABCE1 mutant with a myc-tagged version of the protein lacking the N-terminal extension (myc-TgABCE1 Comp ΔNt, **Fig. S4**). The truncated myc-tagged version was found to localize to the cytoplasm and was expressed constitutively when the ATc-regulatable copy of TgABCE1 was depleted by ATc (**Fig. 3C, D**). Interestingly, parasites expressing only the N-terminally truncated copy were displaying an intermediate phenotype still able to generate some plaques (**Fig. 3E, F**), but were still accumulating some LDs (**Fig. 3G, H, I**). These results indicate that the N-terminal extension is important, but not strictly essential for TgABCE1 function. It could be potentially contributing to the interaction with TgHCF101 (and thus to Fe-S cluster acquisition), or to post-maturation processes more directly linked to the role of TgABCE1 in the parasite.

**Figure 3.**
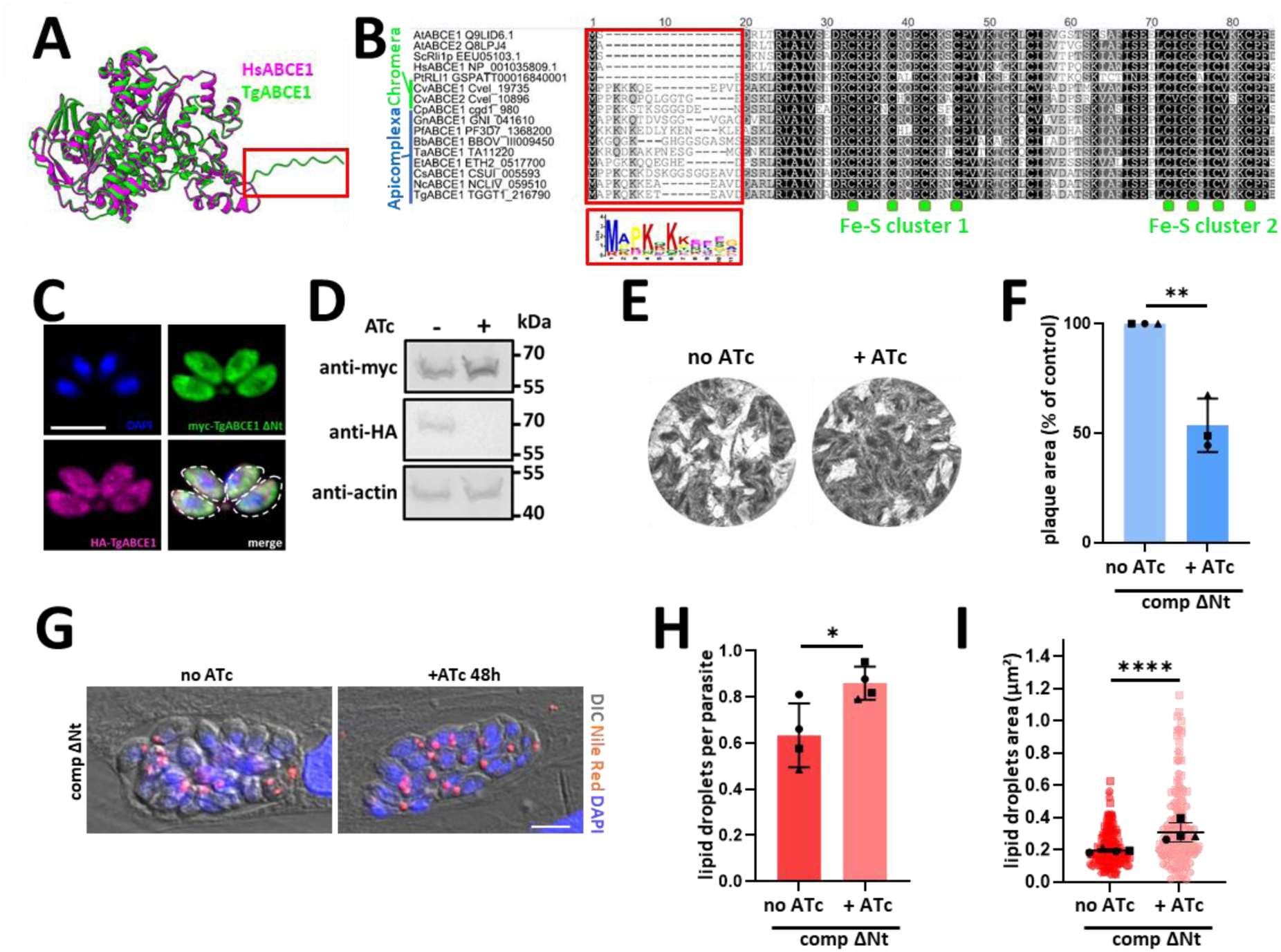
Apicomplexa ABCE1 homologs possess a N-terminal extension that is important, but not essential for function. **A.** Superposition of the AlphaFold-predicted TgABCE1 structure (green) and its *Homo sapiens* homolog HsABCE1 (magenta, protein data bank entry <u>7A09</u>). The red box highlights an N-terminal region only present in TgABCE1. **B.** Amino acid sequence alignment of ABCE1 homologs from *Arabidopsis Thaliana* (GenBank entries: Q8LPJ4, Q9LID6.1), *H. sapiens* (GenBank entry: NP_001035809.1), *Saccharomyces cervisiae* (GenBank entry: EEU05103.1), *Paramecium tetraurelia* (GenBank entry: GSPATT00016840001), *Chromera Velia CCMP2878* (VeupathDB entries: cvel_19735 and cvel_10896), *Cryptosporidium parvum Iowa II* (VeupathDB entry: cgd1_980), *Gregarina niphandrodes* (VeupathDB entry: GNI_041610), *Plasmodium falciparum 3D7* (PF3D7_1368200), *Babesia bovis T2Bo* (BBOV_III009450), *Theileria annulata Ankara* (VeupathDB entry: TA11220), *Eimeria tenella Houghton 2021* (VeupathDB entry: ETH2_0517700), *Cystoisospora suis Wien I* (VeupathDB entry: CSUI_005593), *Neospora caninum Liverpool* (VeupathDB entry: NCLIV_059510),*T. gondii* (VeupathDB entry: TGGT1_216790). The green squares indicate the putative cysteines involved in Fe-S cluster coordination. The red box highlights the specificity of the N-terminal region found only in Apicomplexa containing a potential lysine-rich motif identified by the MEME suite. **C.** IFA confirms the cytoplasmic localization of the myc-tagged extra copy of TgABCE1 in which the N-terminal part has been deleted (myc-TgABCE1 comp ΔNt). Parasite shapes are outlined. DNA was stained with DAPI. Scale bar = 5 μm. **D.** Immunoblot analysis showing expression of the myc-tagged N-terminally-deleted extra copy of TgABCE1 and the regulated depletion of the ATc-regulated HA-tagged endogenous copy TgABCE1. Actin was used as a loading control. **E.** Plaque assay showing partial restoration of growth by the N-terminally-deleted extra copy of TgABCE1 upon depletion of the ATc-regulated copy. Plaque assay was carried out by infecting a monolayer of HFFs with the cKD HA-TgABCE1 comp ΔNt cell line for 7 days in the presence or absence of ATc. **F.** Quantification of plaque area observed in (**E**). Results are expressed as a percentage of the lysed area relative to the control incubated without ATc (set as 100% for reference). Values represented are mean ± SD from *n* = 3 independent biological replicates, ns: not significant (*p*-value >0.05), *p*-values from Student’s t-test. **G.** Representative images of vacuoles containing parasites grown continuously in the presence of ATc for 48 h and stained by Nile Red (orange) for lipid droplets. DIC: differential interference contrast. DNA was stained with DAPI. Scale bar = 5 µm. **H, I.** Quantification of LD area and number, respectively; 100 parasites were analyzed per condition. The cKD HA-TgABCE1 comp ΔNt parasites were grown with ATc for 48 h. Values are represented as the mean ± SD of *n* = 3 independent biological replicates (different symbols represent different series); **** *p*-value ≤0.0001, Student’s t-test.

### TgABCE1 is a key regulator of parasite protein synthesis

ABCE1 is a highly conserved protein in eukaryotes and Archaea^33^ with a major role in regulating translation. The TgHCF101 mutant showed an impact on the translation rate, which was potentially linked to a decrease in TgABCE1 abundance^21^, but the direct implication of the latter in the regulation of parasite protein synthesis remained to be demonstrated. Puromycin is a naturally occurring aminonucleoside mimicking the 3′ end of aminoacylated tRNAs that inhibits protein synthesis, but can also be used as a probe for measuring protein elongation rates as it is incorporated into elongating nascent chains^34^. We assessed puromycin incorporation by immunoblot and IFA, and observed that down-regulation of TgABCE1 expression with ATc led to a strong decrease in puromycin labeling in cKD HA-TgABCE1 parasites, similar to the use of translation inhibitor cycloheximide (**Fig. 4A, B, C**).

**Figure 4.**
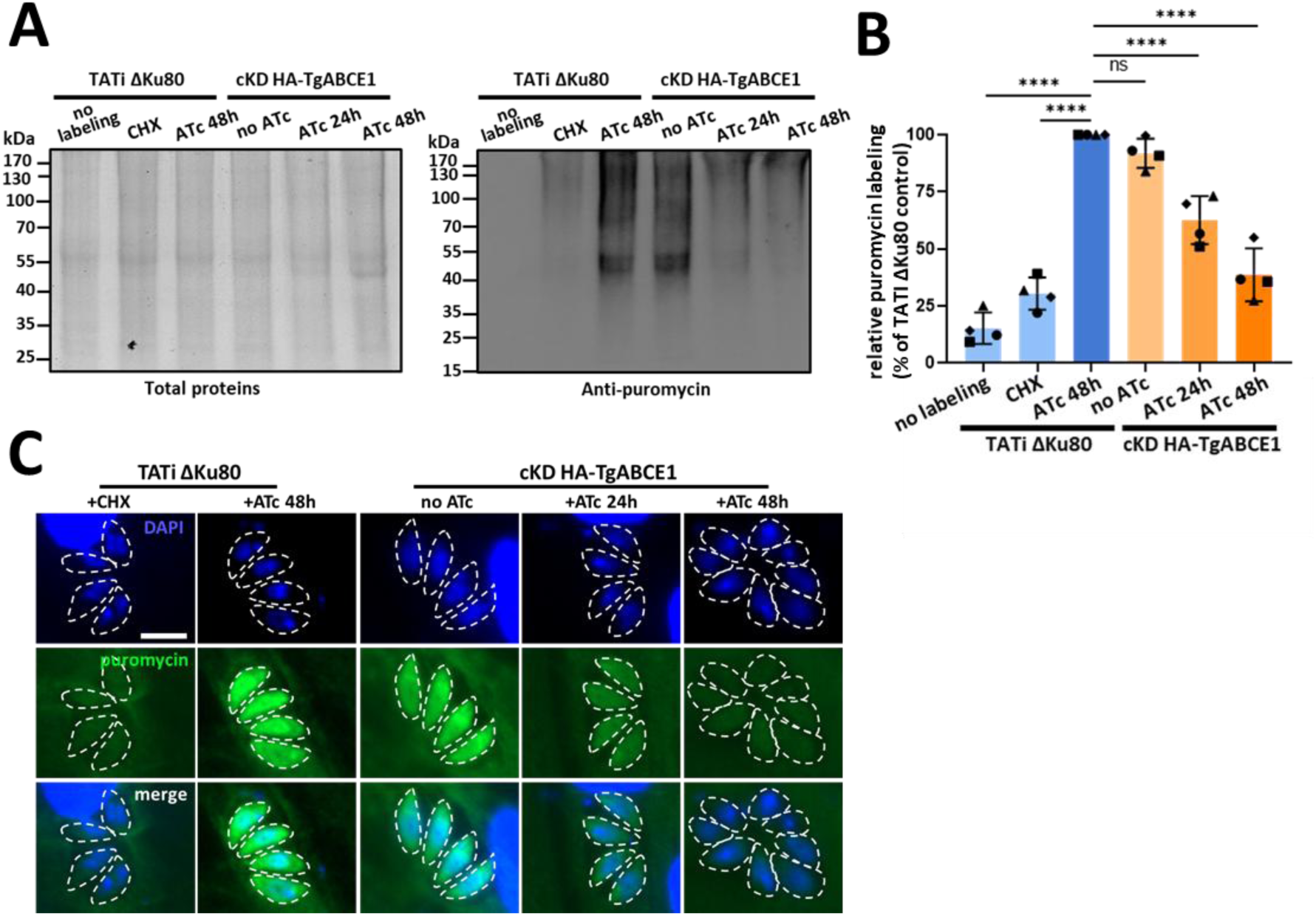
TgABCE1 has a conserved role in protein synthesis. **A.** Immunoblot analysis of puromycin incorporation in the parental (TATi ΔKu80) and cKD HA-TgABCE1 cell lines, untreated or treated with ATc for 48 h. The parasites were treated with 100µg/ml puromycin for 15min. TATi ΔKu80 treated with cycloheximide (CHX) at 100µg/ml was used as a control for translation inhibition. The puromycin signal was detected with anti-puromycin antibody (right), and total protein content was visualized by stain-Free imaging technology (left). **B.** Variation of puromycin incorporation in the different conditions was quantified by band densitometry and normalized on the total protein content of each respective lane. Puromycin labeling is presented as a percentage relative to the control TATi ΔKu80 +ATc, set as 100% for each biological replicate. Values are represented as the mean and SD of 4 independent biological replicates; ns : not significant (*p*-value ≥0.05), **** *p*-value ≤0.0001. *p*-values from one-way ANOVA with Dunnett’s multiple comparison test. **C.** Puromycin incorporation visualized by immunofluorescence in parental (TATi ΔKu80) and cKD HA-TgABCE1 parasites treated or not with ATc for 48 h. The parasites were treated with puromycin for 15 min. TATi ΔKu80 treated with CHX was used as a control for translation inhibition. The puromycin signal was detected with anti-puromycin antibody (green) and DNA is stained with DAPI. Parasite shapes are outlined. Scale bar = 5 μm.

Puromycin-based assays, which are based on a short pulse labeling, allow assessing protein elongation rates, but to get further insights into the long-term impact of TgABCE1 depletion on protein synthesis, we next used L-azidohomoalanine (AHA), a clickable analog of methionine that can be incorporated into newly synthesized proteins^35^. We performed AHA incorporation for up to 6 h, in conditions that we have previously shown to be suitable for protein labeling without affecting parasite viability^36^. Using a clickable fluorophore, we detected AHA labeling by microscopic imaging in both host cells and parasites (with stronger labeling in the latter) after 6 h of incubation. Labeling was largely prevented by cycloheximide treatment and was greatly reduced in parasites upon ATc-mediated downregulation of TgABCE1 expression (**Fig. 5A**). We combined clickable AHA labeling with affinity purification and mass spectrometry to obtain a quantitative and qualitative snapshot of proteins synthesized in the presence or absence of TgABCE1, as well as following incubation with cycloheximide (as a translation inhibitor control). Parasites were incubated for 6 h with AHA under various treatment conditions, including the use of cycloheximide, or incubation with ATc to deplete TgABCE1. Newly synthesized polypeptides incorporating AHA were then selectively purified by click chemistry on an alkyne agarose column, followed by on-column trypsin digestion and analysis by quantitative mass spectrometry (**Fig. 5B**). This approach provided an overview of the proteins synthesized during the 6 h AHA incubation period, along with the abundance of their corresponding peptides (**Table S1**). When peptide abundance was plotted to compare the different conditions (**Fig. 5C**), it showed that the majority of proteins had more peptides detected in the TgABCE1-expressing condition than when it was depleted by the use of ATc or when cycloheximide was added. In contrast, the addition of ATc or cycloheximide had very similar effects on peptide abundance across the detected proteome, with the notable exception of TgABCE1, for which even fewer peptides were detected upon ATc treatment, thereby validating the efficient downregulation of its expression.

**Figure 5.**
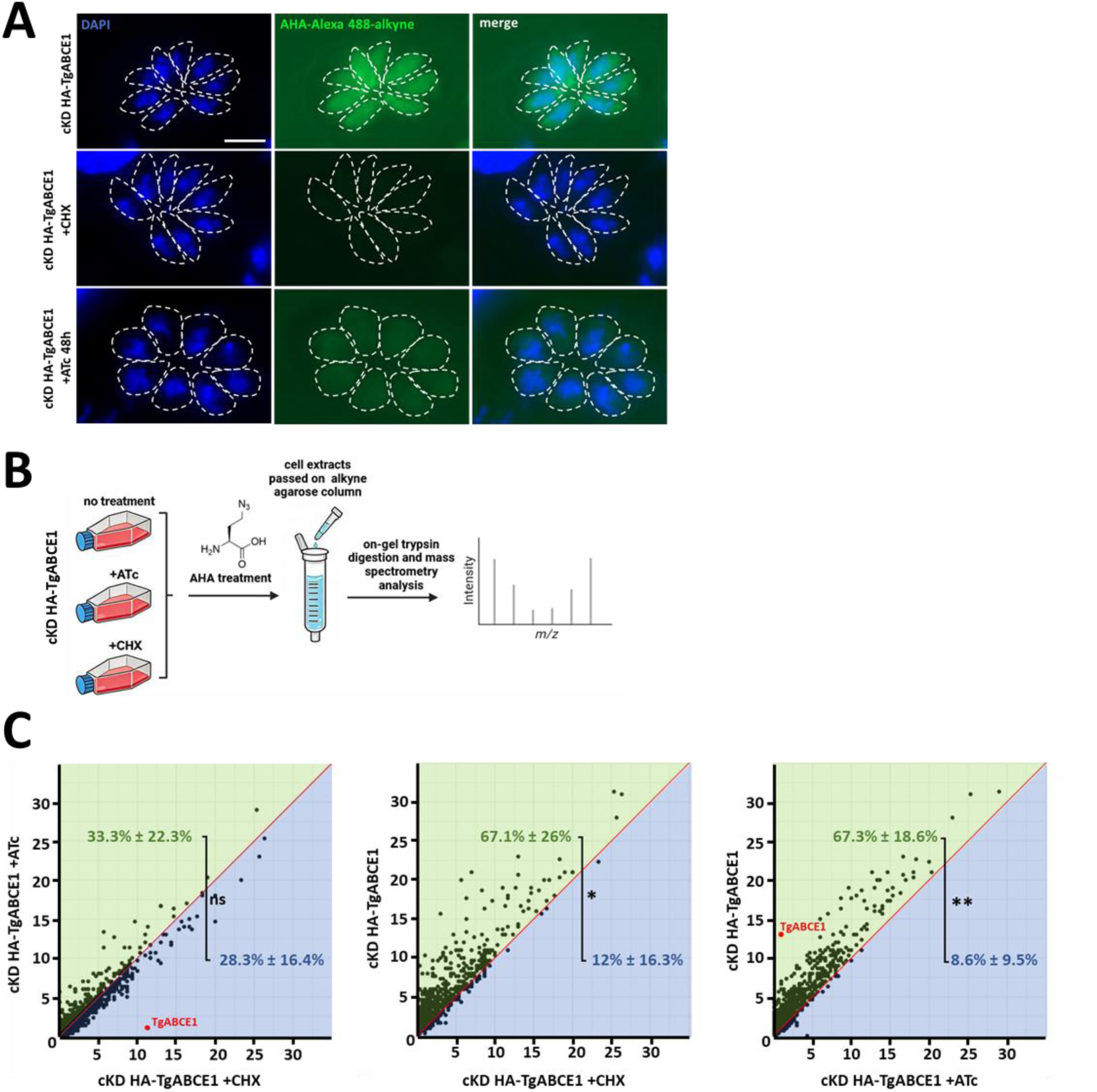
TgABCE1 depletion leads to a general drop in protein synthesis. **A.** Immunofluorescence analysis of L-Azidohomoalanine (AHA, a ‘clickable’ methionine analog) labeling in the cKD HA-TgABCE1 cell line, either untreated or treated with ATc for 48 h. The cKD HA-TgABCE1 line treated with CHX for 6 h served as a control for protein synthesis inhibition. The parasites were treated for 30 min in methionine-free DMEM and for 6 h with 25 µM AHA in methionine-free DMEM. Subsequently, the incorporation of AHA into newly synthesized proteins was directly detected via a ‘click’ chemistry reaction between the azide group of AHA and the Alexa-488-conjugated fluorescent alkyne. Parasite shapes are outlined and the DNA was stained with DAPI. Scale bar = 5 μm. **B.** Schematic outline of the AHA labeling protocol for mass spectrometry-based peptide quantification: cKD HA-TgABCE1 parasites were treated or not with ATc for 48 h, or with CHX. After 42 h in the presence or absence of ATc, the intracellular parasites were placed in a methionine-free medium for 30 min., then in methionine-free DMEM medium enriched with 25 µM of AHA for 6 h. The parasites were then lysed, and the ‘click’ reaction was carried out on an alkyne-agarose column from the click-It kit. The proteins retained on the column were eluted, digested, and analysed by mass spectrometry. **C.** Graph showing the average number of peptides identified by mass spectrometry for a given protein under each condition. The red diagonal line indicates the same number of peptides for the same protein under each condition. Mean percentage of proteins with higher peptide number is displayed for each condition (±standard deviation, *n* = 3). ns, not significant (*p*-value >0.05), * *p*-value ≤0.05, ** *p*-value ≤0.01, Student’s t-test.

Overall, our data confirm that TgABCE1 plays a pivotal role in regulating global protein synthesis in the parasite.

### Blocking protein synthesis and parasite growth leads to an imbalance in lipid homeostasis

We previously observed the accumulation of LDs in conditions of iron starvation^27^, but also in mutants related to the CIA pathway^21^, such as the TgNAR1 protein that connects the early and late steps of the pathway, and TgHCF101 that potentially transfers Fe-S clusters to TgABCE1 (**Fig. S5**). In contrast, there was no pronounced LDs accumulation when blocking the Fe-S assembly pathways hosted by the parasite’s endosymbiotic organelles like the SUF or ISC pathway^21^ (**Fig. S5**). Given the phenotypic convergence towards Fe-S protein TgABCE1, and the role of the latter in protein synthesis, we decided to check a potential impact of translation inhibitors on LDs. We used either puromycin or cycloheximide, incubated for either 6 h or 16 h, and quantified LDs (**Fig. 6A, B, C, D**). Both inhibitors led to an increase in LD number and size and, strikingly, also seemed to induce LD accumulation in host cells (**Fig. 6A, B**).

**Figure 6.**
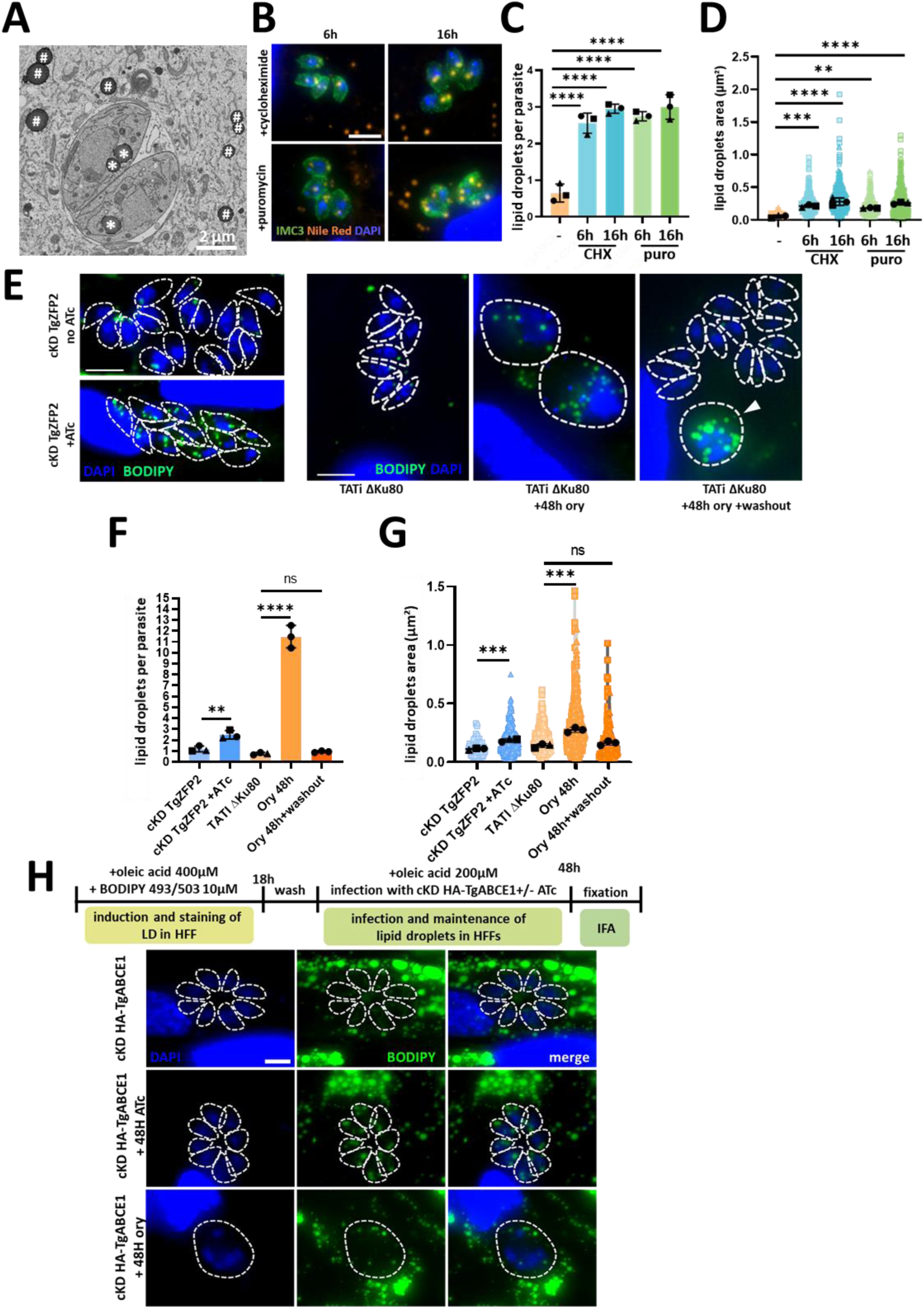
Inhibition of protein synthesis and block of cell division both lead to an accumulation of lipid droplets. **A.** Electron micrograph of intracellular parasites treated with cycloheximide (CHX) for 6 h. Asterisks and square signs denote parasite and host lipid droplets (LDs), respectively. Pixel size: 10 nm. **B.** Immunofluorescence microscopy imaging of parasites untreated or treated for 6 h and 16 h with translation inhibitors cycloheximide and puromycin. LDs accumulate in both parasites and host cells. LDs were stained with Nile Red (orange), parasites were labeled with an anti-IMC3 antibody (green), and DNA was stained with DAPI. Scale bar = 5 μm. For this experiment, puromycin and CHX were used at 100 µg/ml. **C, D.** Quantification of lipid droplet number and area in cKD HA-TgABCE1 parasites grown in the absence or in the presence of CHX and puromycin for 6 h or 16 h. A total of 100 parasites were analyzed per condition. Values are represented as the mean ±standard deviation of *n* = 3 independent biological replicates (different symbols represent different series); ns: not significant (*p*-value >0.05), **: *p*-value ≤0.01, ****: *p*-value ≤0.0001. *p*-value*s* from one-way ANOVA with Dunnett’s multiple comparison test. CHX: cycloheximide, puro: puromycin, SD: standard deviation. **E.** Immunofluorescence imaging of parasites from the cKD HA-TgZFP2 cell cycle mutant grown in the absence or presence of ATc 48 h, in which the ZFP2 depletion induces an accumulation of LDs (left). Immunofluorescence imaging of parasites pre-cultured for 24 h and then incubated for 48 h with the cell cycle inhibitor oryzalin (ory, 2.5 µM) results in the accumulation of LDs (middle). Ory was then washed out and parasites were left to recover for an extra 48 h : while some vacuoles remained blocked (arrowhead), parasites resuming cell division displayed less LDs (right). LDs were stained with BODIPY 493/503 (green), parasites are outlined, and DNA was stained with DAPI (blue). Scale bar = 5 μm. **F, G.** Quantification of LD number and area in parasites depleted of TgZFP2, or treated with ory, with or without washout. 100 parasites were analyzed per condition. Values are represented as the mean ± SD of *n* = 3 independent biological replicates (different symbols represent different replicates); **: *p*-value ≤0.01, ***: *p*-value ≤0.001, ****: *p*-value ≤0.0001, from Student’s t-test. **H.** Outline of the host cell lipid scavenging protocol: before infection with the parasites, HFF cells were treated for 18h with BODIPY 493/503 and oleic acid (OA, 0.4 mM) to stimulate the accumulation of host LDs. After washing, the HFFs were infected with cKD HA-TgABCE1 in the absence or presence of ATc for 48 h or treated with 2.5 µM ory. OA concentration was maintained at 0.2 mM to sustain host lipid droplets production. Immunofluorescence imaging shows, in both cases, the BODIPY 493/503 signal incorporated in the parasites’ LDs, illustrating scavenging from the host.

We revisited our proteomic dataset (**Table S1**) to identify the few proteins that were specifically increased in abundance and shared between the TgABCE1-depleted and cycloheximide-treated conditions. We reasoned that some lipid-modulating protein factors might be upregulated under these conditions. However, we did not identify any obvious candidate that may be potentially LD-inducing. LD accumulation may also be a consequence of a decrease in abundance of protein factors (some involved in lipid turnover for instance), but as a majority of proteins decreased in abundance in TgABCE1-depleted and cycloheximide-treated conditions, it was also difficult to point to specific lipid-related candidates. Instead of specific factors being upregulated, we then thought that LD accumulation could be a more general consequence of protein synthesis block. Parasites actively scavenge resources (including lipids^37^) from their host, and although they likely undergo a relatively rapid block in cell division upon depletion of TgABCE1 (**Fig. 1F**, possibly due to inhibition of protein synthesis, which is essential for cell-cycle progression^38^), they may still be able to take up lipids from the host before dying. To test this hypothesis, we quantified LDs in a mutant of the zinc finger protein TgZFP2, whose depletion blocks cell cycle progression^39^, as well as after treatment with oryzalin, a microtubule-disrupting compound that leads to a similar effect^40^ (**Fig. 6E, F, G**). Strikingly, both genetic and chemical disruption of the cell division process led to an accumulation of LDs. Moreover, upon oryzalin washout, some blocked parasites were able to resume division (66%±12%, *n* = 3), and in those LDs decreased considerably in number and size, indicating a dynamic and reversible process (**Fig. 6E, F, G**).

We next wanted to assess whether parasites could still scavenge neutral lipids originating from host LDs upon TgABCE1 depletion or cell cycle arrest. As described previously^41^, we thus pre-labelled host LDs with the non-metabolizable lipid dye BODIPY 493/503 in the presence of oleic acid, resulting in a high number of fluorescent LDs in the host. Then, after washing, we infected these host cells with parasites in which we either depleted TgABCE1 or blocked cell cycle progression with oryzalin (**Fig. 6H**). Fluorescent LDs were observed in both TgABCE1-depleted or oryzalin-treated parasites, demonstrating that they could incorporate scavenged neutral lipids from the host.

Our results thus show that the inhibition of protein synthesis resulting from TgABCE1 depletion may cause a progressive arrest in cell division, but that the parasite can still scavenge host lipids, leading to an accumulation of LDs.

### The interactome of TgABCE1 identifies a new protein involved in the parasite stress response

In order to get further insights into the potential involvement of TgABCE1 in larger biological processes, we performed co-immunoprecipitation experiments, followed by mass spectrometry-based identification of its putative partners (**Fig. 7A**). We uncovered a limited number of candidates that included TgHCF101, the *T. gondii* homolog of MAK16 (TGGT1_249950 entry in the https://toxodb.org database), and an uncharacterized protein (TGGT1_218950). ABCE1 has been found in other eukaryotes to associate with ribosomal subunits and translation initiation factors^9,10^, but we did not recover these components with sufficient statistical significance with our analysis (**Table S2**), possibly reflecting the transient nature of these interactions.

**Figure 7.**
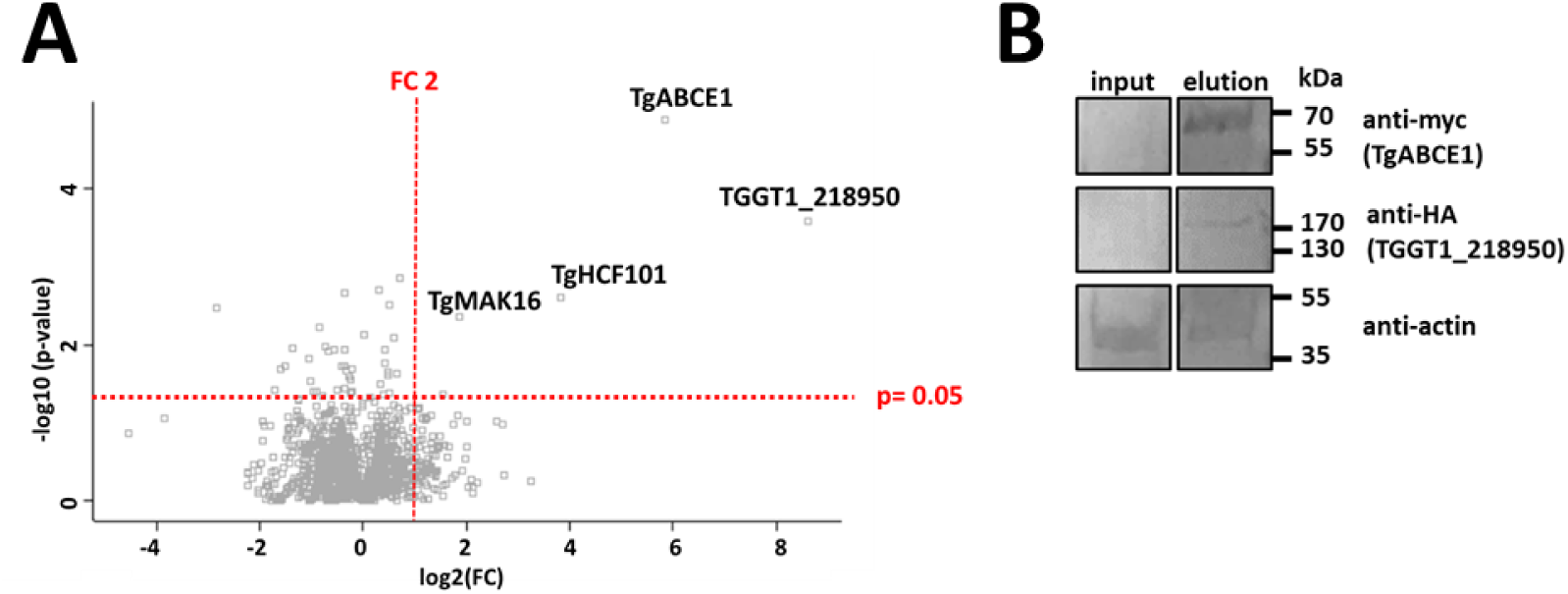
Identification of potential TgABCE1 partners. **A.** Volcano plot showing differential expression of TgABCE1 and co-immunoprecipitated proteins in the cKD HA-TgABCE1 and TATi ΔKu80 cell lines, as analyzed by quantitative proteomic. Cut-offs were set at ≥2-fold change and *p*-value ≤0.05. More abundant proteins are annotated on the plot. **B.** Immunoblot analysis of the TGGT1_218950-mAID-HA_3_ myc-TgABCE1 myc-cell line after myc-tag affinity purification, revealing an enrichment of both myc-tagged TgABCE1 and HA-tagged TGGT1_218950 on the column versus the initial input. Actin, used as a control, although detected in the elution fraction, is not enriched compared with the input.

Nevertheless, the identification of TgHCF101 among the co-immunoprecipitated proteins validated the quality of our experimental results, as TgABCE1 was initially found to associate with TgHCF101 in the reverse co-immunoprecipitation, and both proteins were further confirmed to interact directly by yeast two-hybrid analysis^21^. TGGT1_249950 is the *T. gondii* equivalent of MAK16, a protein that plays an early role in the maturation of the large ribosomal subunit of eukaryotes^42^. MAK16 typically functions in the nucleolus, where preribosomes are being assembled, and while we did not manage to tag TgMAK16, which would have allowed us to verify its localization (or its binding to TgABCE1), it seems to have conserved consensus nuclear localization sequences^43^ (**Fig. S6A**). The yeast ABCE1 homolog has also been functionally linked to ribosome assembly^15^and suggested to shuttle between the nucleus and the cytosol, which is not obvious for TgABCE1 (**Fig. 1B**). Thus, while TgABCE1 and TgMAK16 are both linked to ribosomal function, it is unclear if they could interact directly in the context of ribosomal assembly, as they may localize to different compartments. However, interestingly, MAK16 has been found to co-immunoprecipitate with members of the CIA complex in human cells^44^, and was very recently confirmed to be a Fe-S cluster protein^45^. Importantly, the cysteines potentially coordinating the Fe-S cluster are conserved in TgMAK16 (**Fig. S6A**), and although it was not initially identified in the repertoire of co-immunoprecipitated partners of TgHCF101^21^, TgMAK16 and TgABCE1 may both be its client proteins.

One last candidate for interaction was a large uncharacterized protein (TGGT1_218950, predicted to be about 126 kDa) with no particular primary sequence homology to any known protein, with as its only identifiable feature the presence of armadillo/HEAT repeat-type folds^46^. AlphaFold prediction, albeit mostly displaying low confidence scores, confirmed that TGGT1_218950 is essentially composed of α-helical folds (**Fig. S6B**). Further analysis with HHpred^47^ (allowing remote homology detection through comparison of hidden Markov models) found homology with armadillo repeat proteins, including GCN1 (**Fig. 6C**). Of course, this may just be because GCN1 is composed almost entirely of α-helical HEAT repeats^48^. Moreover, another *T. gondii* protein (TGGT1_231480), although not yet fully characterized, is more likely to be the true GCN1 homolog of the parasite^49^. However, although remote and partial, this structural homology caught our attention because GCN1 is involved in ribosome quality control^50^, which may functionally link it to ABCE1^51^.

We thus generated a conditional knock-down cell line for TGGT1_218950 using the auxin-inducible degron (AID) system in the RH ΔKu80 TIR1 strain^52^ background (**Fig. S7**). This allowed the addition of a C-terminal AID module, along with a triple HA tag to TGGT1_218950. We next expressed in that background a myc-tagged copy of TgABCE1 (**Fig. S8A, B**). IFA on this cell line showed that TGGT1_218950 seems to localize to the cytosol, like TgABCE1 (**Fig. S8C**). We then performed co-immunoprecipitation experiments to investigate further the potential interaction between TgABCE1 and TGGT1_218950. Immunoblot analysis confirmed the association of HA-tagged TGGT1_218950 with immunoprecipitated myc-tagged TgABCE1 (**Fig. 7B**), although the reverse immunoprecipitation was unfortunately inconclusive, perhaps due to a low level of TGGT1_218950 expression. Immunolabeling verified that TGGT1_218950 seemed to be expressed at low basal levels indeed, but also confirmed it was efficiently down-regulated in a few hours thanks to the AID strategy (**Fig. 8A, B**). Depletion of the protein led to an absence of plaques, suggesting its essentiality for parasite growth (**Fig. 8C**). TGGT1_218950 was also shown to be essential for intracellular parasite replication (**Fig. 8D**), albeit with a less pronounced effect than TgABCE1 depletion (**Fig. 1E**). LDs were found to increase in both number and size upon TGGT1_218950 depletion (**Fig. 8E, F, G**), although again in a less pronounced manner compared with those observed in absence of TgABCE1 (**Fig. 1I**). Moreover, in sharp contrast with TgABCE1, the absence of TGGT1_218950 did not have a strong immediate impact on protein synthesis as assessed by puromycin labeling (**Fig. S9**). Yet, as GCN1-related proteins are typically activators of the GCN2 kinase, involved in the response to ribosomal collision in stress conditions^53^, we sought to evaluate the potential consequences of TGGT1_218950 depletion in the survival of extracellular parasites, for which TgGCN2 plays a key role^54^ (**Fig. 8H, I**). We found that TGGT1_218950 is important for maintaining the viability of extracellular tachyzoites but, interestingly, not TgABCE1.

**Figure 8.**
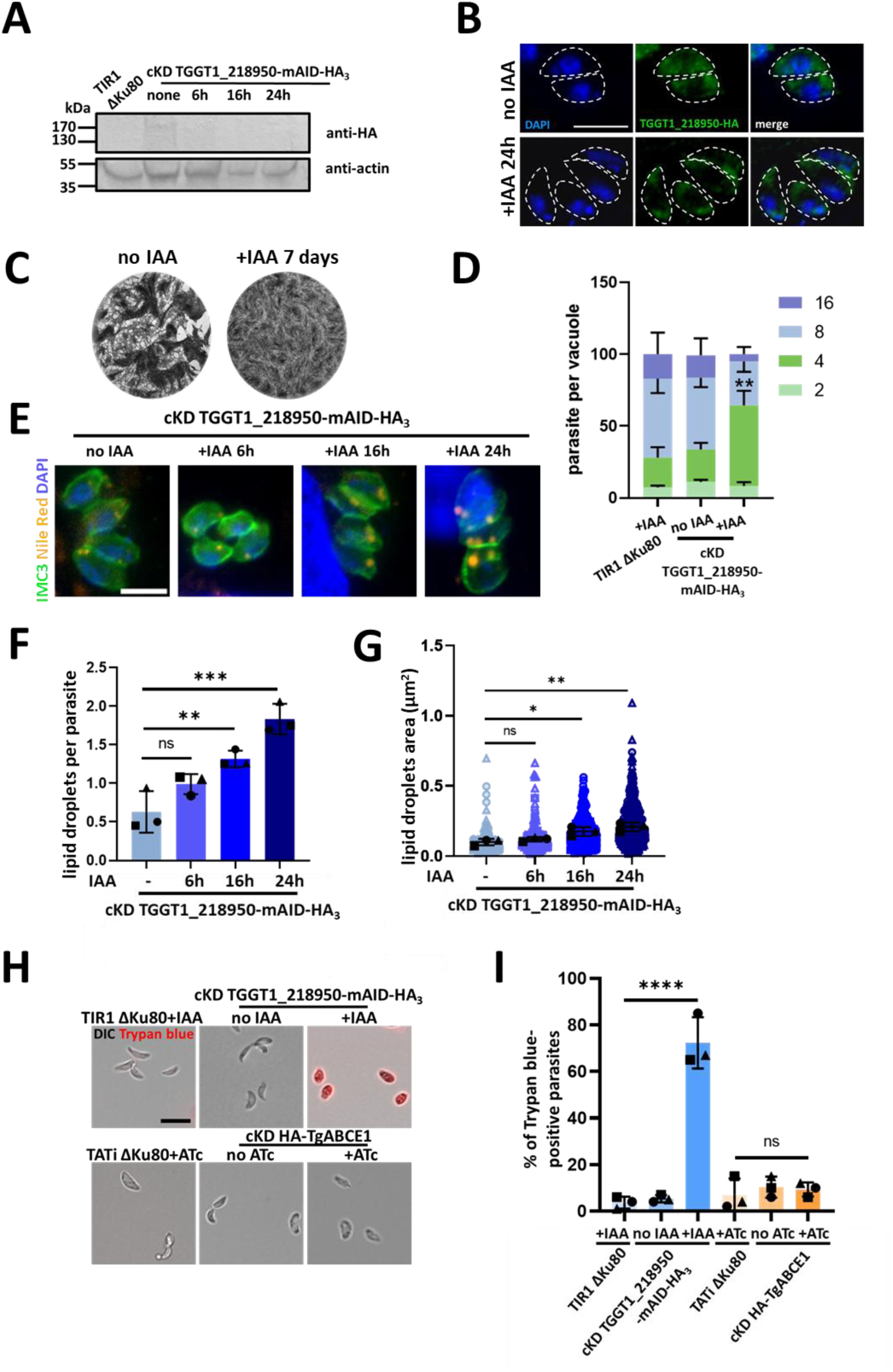
Phenotypic characterization of TGGT1_218950. **A.** Immunoblot analysis of the cKD TGGT1_218950-mAID-HA_3_ mutant showing efficient tagging and down-regulation of TGGT1_218950 starting at 6 h of treatment with indole-3-acetic acid (IAA). Actin was used as a loading control. **B.** Immunofluorescence assay showing cytosolic localization of the HA-tagged TGGT1_218950 protein, and depletion of the protein after IAA addition. Parasites are outlined and DNA was stained with 4′,6-diamidino-2-phenylindole (DAPI). Scale bar = 5 μm. **C.** Plaque assays showing that TGGT1_218950 knockdown renders the parasites unable to form plaques in the host cell monolayer. These tests were carried out by infecting a monolayer of HFFs with the cKD TGGT1_218950-mAID-HA_3_ cell line for 7 days in the presence or absence of IAA. **D.** Replication assay of parental (TIR1 ΔKu80) and transgenic cell line (cKD HA-TgABCE1): parasites were pre-cultured for 24 h in the presence or absence of ATc and allowed to invade HFF-coated coverslips for another 24 h in the presence or absence of ATc, for a total of up to 48 h. The number of parasites per vacuole was quantified for each condition and expressed as a percentage; 100 vacuoles were counted for each condition. Values represented are mean ± SD of *n* = 3 independent biological replicates. Statistical significance was determined between each category by Student’s t-test and noted as follows: ** *p*-value ≤0.01. **E.** Immunofluorescence imaging of LD present upon depletion of TGGT1_218950. Parasites are outlined and DNA was stained with 4′,6-diamidino-2-phenylindole (DAPI). Scale bar = 5 μm. **F, G.** Quantification of LD number and area in parasites depleted of TGGT1_218950. 100 parasites were analyzed per condition. Values are represented as the mean ± standard deviation of *n* = 3 independent biological replicates (different symbols represent different replicates); *: *p*-value ≤0.05, **: *p*-value ≤0.01, ***: *p*-value ≤0.001, from Student’s t-test. **H.** Viability of extracellular parasites (previously grown in the presence or absence of ATc for 48 h, or IAA for 24 h) kept in DMEM medium for 6 h, as assessed by their permeability to Trypan blue, which is impermeable to intact membranes and becomes fluorescent when complexed with intracellular proteins. **I.** Quantification of the percentage of Trypan blue-positive parasites exposed to extracellular stress as described in **H**. Values are presented as mean ± standard deviation of *n* = 3 independent biological replicates (different symbols represent different series): ns: not significant (*p*-value ≥0.05), **** *p*-value ≤ 0.0001. *p*-values derived from a one-way ANOVA with Dunnett’s multiple comparison test.

Thus, while TgABCE1 may be interacting with TGGT1_218950 in the context of translational regulation, our results suggest that the latter seems to be more specialized in the stress response.

## Discussion

Fe-S clusters are cofactors for proteins involved in essential housekeeping functions across all kingdoms of life. This is reflected in the predicted Fe-S proteome of *T. gondii*, which was identified through bioinformatic predictions followed by manual curation^22^. Among these candidates, TgABCE1 is arguably one of the most highly conserved Fe-S proteins across eukaryotes and Archaea, where it is essential for viability^16^. Several CIA mutants in *T. gondii* exhibit reduced ABCE1 protein levels, together with a possible decrease in translation rates^21,28,55^, although the function of ABCE1 itself had not previously been directly characterized in the parasites. The present work establishes the conserved function of ABCE1 in *T. gondii*, as protein synthesis rate (assessed by puromycin incorporation assays) and peptide quantification in nascent proteins were found to be strongly reduced upon TgABCE1 depletion (**Figs. 4 and 5**).

However, while ABCE1/Rli1 depletion in eukaryotes typically reduces global protein synthesis, the most prominent defect can vary across organisms and experimental systems. The best-established role for this protein is in ribosome recycling after translation termination, but it has also been linked to translation initiation and ribosome biogenesis, suggesting that these functions may be complementary aspects of a multifunctional factor that helps maintain ribosome homeostasis and translation efficiency^7,6^. We performed co-immunoprecipitation experiments to gain further insight into the precise function of TgABCE1 in protein synthesis, but identified relatively few candidates for interacting partners. One of them, was the *T. gondii* homolog of MAK16, a protein involved in the assembly of the 60S ribosome in the nucleolus^43^. We could not tag TgMAK16 to verify its nuclear localization and, although we found primarily TgABCE1 localized to the cytoplasm (**Fig. 1B**), we cannot exclude that it is also partly present in the nucleus. Early studies actually proposed that the yeast ABCE1 homolog can shuttle between the cytosol and the nucleus where it may play a role in ribosome assembly^15^, which has not been firmly established in other systems since. Yet, the involvement of TgABCE1 in ribosomal assembly together with TgMAK16 remains a possibility. As MAK16 has recently been shown to be a Fe-S protein itself^45^, so another possibility is that it would share its transfer protein with ABCE1 and that they would be present together in a protein complex during their maturation.

Another potential TgABCE1 partner identified through co-immunoprecipitation is the uncharacterized protein TGGT1_218950, which contains no significant sequence or structural similarity to any other protein except for α-helical folds resembling armadillo/HEAT repeats. These motifs are also present in GCN1, a stress-induced regulator of ribosome-mediated signaling^51^. While we have demonstrated the essentiality of TGGT1_218950 for parasite growth and its potential importance in coping with extracellular stress, its phenotype does not fully overlap with that of TgABCE1 (**Fig. 8**). Therefore, further investigation will be required to assess the potential involvement of TGGT1_218950 in the ribosome collision response, or to confirm a potential functional relationship with TgABCE1

Co-immunoprecipitation also confirmed the association of TgABCE1 with its potential Fe-S cluster transfer protein, TgHCF101. ABCE1 typically contains two Fe-S clusters, but while they are delivered from the CIA assembly pathway by specific transfer proteins in opisthokonts^20^, they are absent from several eukaryotic lineages, including Apicomplexa where HCF101 is playing that role instead^21^. Although Archaea have ABCE1 homologs, they lack a CIA pathway, relying instead on more rudimentary and ancient assembly systems^56^, thus suggesting that cluster transfer to ABCE1 likely specialized independently in different lineages. The HCF101/ABCE1 interaction being essential and specific to Apicomplexa, its disruption may have some therapeutic potential. It would thus be important to determine the molecular determinants of this interaction to design some specific inhibitors. We showed, for example, that the Apicomplexa-specific N-terminal extension of TgABCE1 is important, although not essential, for its function (**Fig. 3**). However, further investigation, including AlphaFold-guided identification and mutational analysis of potential key residues, will be required.

This work validates TgABCE1 as a client protein of TgHCF101, not only because their interaction has been confirmed, but also because depletion of TgABCE1 largely phenocopies that of TgHCF101 at the ultrastructural level (cell division defects and accumulation of LDs, **Fig. 1**). Of course, these phenotypic manifestations could be due to the perturbation of other proteins involved in regulating translation or cell division. Thus, while TgABCE1 is the Fe-S protein with the strongest evidence of interaction with TgHCF101, it is also possible that the latter acts as a transfer protein to other proteins involved in these cellular processes. It was recently shown that global iron depletion in the parasite leads to translational repression prior to a decrease in TgABCE1 abundance^24^, suggesting that it may not be the main iron-dependent driving factor for regulating protein synthesis. It is certainly not the only one, as the predicted Fe-S proteome includes several proteins involved in translation^22^, and ribosomal assembly factors like TgMAK16 (a potential Fe-S protein^45^, **Fig. S6**) are likely contributing to iron-dependent protein synthesis. Yet, protein abundance is not a perfect readout for protein functionality (some Fe-S proteins retain some stability in spite of the absence of their Fe-S clusters), and we have demonstrated here the strong dependence of TgABCE1 on its Fe-S clusters for function (**Fig. 2**), hinting that it would be highly sensitive to iron deprivation. Moreover, our work shows that depletion of TgABCE1 leads to a strong and direct impact on protein synthesis. Interestingly, while translational repression has been shown to accompany differentiation into bradyzoites^57,4,58,59^, we did not observe extensive induction of stage conversion upon TgABCE1 depletion (**Fig. S2A**).

One striking phenotype we observed is LD accumulation, which occurred not only upon depletion of the CIA-dependent TgABCE1, but also under iron starvation and in CIA pathway mutants, whereas it was not observed following disruption of the SUF or ISC pathways^21^ (**Fig. S5**). These results hinted for a convergence toward the translation-regulation function of Fe-S protein TgABCE1 and, more generally, on its potential impact on cell growth. Interestingly, acute iron depletion has been shown to induce LD accumulation in mammalian cells^60^, and so does the use of several translation inhibitors^61^ (**Fig. 6A, B**). One possibility is that the inhibition of protein synthesis may suppress lipid turnover by impacting lipid hydrolases that could have relatively short half-lives, while proteins involved in lipid synthesis or acquisition may be more long-lived. In addition, protein synthesis is one of the most energy consuming process in eukaryotic cells, estimated to consume ∼30% of cellular ATP in mammalian cells for instance^62^. Thus, another possibility is that blocking this process could lead to a sharp energetic imbalance, leading to an excess of LDs if they would be used routinely as an ATP source. However, in tachyzoites the primary source of ATP production comes from glycolysis and by catabolizing glucose and glutamine via an oxidative tricarboxylic acid (TCA) cycle^63^. Thus, contrarily to other developmental stages, they likely do not extensively rely on fatty acid breakdown through β-oxidation to fuel the TCA cycle through the production of acetyl-CoA^64^. Interestingly, recent evidence indicates that drug-induced cell cycle arrest also promotes LD formation in several mammalian cell types^65^ and ABCE1 was shown to be essential for S phase progression in human cells^66^, like it appears to be for *T. gondii* (**Fig. 1F**), suggesting that the underlying mechanism we observed may be broadly conserved across eukaryotes. Clearly, cell-cycle progression is highly dependent on continuous protein synthesis, as some regulators are short-lived and must be constantly replenished^67^. In *T. gondii*, inhibition of protein synthesis rapidly blocks parasite replication, consistent with the requirement for continuous synthesis of cell-cycle regulators to sustain progression through endodyogeny^68,69^. Therefore, we propose that inhibition of protein synthesis, either by drug treatment or TgABCE1 depletion, leads to an arrest in parasite development, which appears to be a primary trigger for LD accumulation. Consistent with this hypothesis, we show that both genetic and pharmacological blockade of parasite cell division leads to increased LD accumulation. Inside host cells, *T. gondii* actively scavenges lipids, especially from host LDs, and uses them to support intracellular growth and membrane biogenesis^41^. Our experimental evidence further suggests that while they are progressively blocked in cell division, parasites retain the capacity to scavenge lipids from the host (**Fig. 6**), which may be subsequently stored in the parasites’ LDs because they can no longer be incorporated into developing daughter cells.

In summary, our study highlights the importance of tight regulation of protein synthesis during the intracellular development of *T. gondii*, and our data emphasize the pivotal role played by the Fe-S protein TgABCE1 in this process. Despite its importance, as TgABCE1 is highly conserved between the parasite and its mammalian host it may not itself represent a suitable drug target for antiparasitic strategies. However, future work should instead focus on further characterizing its interaction with the Fe-S cluster transfer protein TgHCF101, which is absent from the host and may therefore provide a more promising avenue for drug design.

## Materials and methods

### Cell culture

*T. gondii* ME49 (ATCC 50840, type II), RH (ATCC 50174, type I) tachyzoites and derived transgenic cell lines generated in this study were routinely maintained through passages in a monolayer of human foreskin fibroblasts (HFFs, ATCC CRL-1634). HFFs and parasites were cultured in standard Dulbecco’s Modified Eagle Medium (DMEM, Gibco) supplemented with 5% decomplemented fetal bovine serum (FBS, Biosera), 2 mM L-glutamine (Gibco), 100 U/ml penicillin (Gibco), and 100 μg/ml streptomycin (Gibco), in a controlled atmosphere at 37°C and with 5% CO_2_.

Transgenic cell lines were generated either in the RH TATi ΔKu80^70^ genetic background, which lacks the *Ku80* gene to favor homologous recombination and expresses the TATi transactivator required for the TetOff system, or the RH ΔKu80 TIR1 strain^52^, which expresses the TIR-1 receptor for the auxin-inducible degron system (miniAID).

For the positive selection of transgenic parasites carrying resistance cassettes, either for the expression of dihydrofolate reductase thymidylate synthase (DHFR-TS), chloramphenicol acetyltransferase (CAT), or hypoxanthine-xanthine-guanine phosphoribosyl transferase (HXGPRT), they were cultured with 1 μM pyrimethamine (SML3579, Sigma-Aldrich), 20 μM chloramphenicol (C0378, Sigma-Aldrich) or 25 μg/mL mycophenolic acid and 50 μg/mL xanthine, respectively.

Conditional depletion in the TetOff conditional knock-down lines was achieved by incubation with 0.5 μg/ml anhydrotetracycline (ATc, 37919, Fluka) for the indicated duration of the assay. In the case of the knock-down in the TIR1-mAID system, 0.5 mM of auxin indole-3-acetic acid (IAA, Sigma-Aldrich) was added to the medium.

### Generating a conditional TgABCE1 knock-down cell line

To generate the construct for the tetracycline-regulated conditional depletion of TgABCE1 and add an N-terminal HA tag, we amplified by PCR a 756 bp fragment corresponding to the 5′ coding region of TgABCE1 (TGGT1_216790 entry in the https://toxodb.org database) using primers ML6175 and ML6176 (primers used in this study are listed in **Table S3**). After digestion with BglII and NotI, it was inserted into the DHFR-TetO7Sag4-HA plasmid^71^. The TATi ΔKu80 cell line was transfected by electroporation with 80 μg of the linearized DHFR-TetO7Sag4-HA-TgABCE1 plasmid with ASiSI enzyme. Transgenic parasites, designated cKD HA-TgABCE1, were selected using pyrimethamine and then cloned by limiting dilution. Positive clones were verified by PCR with primers ML1771 and ML6180.

### Generating complemented cell lines

All complemented cell lines were generated by CRISPR-Cas9-mediated homology-directed recombination at the uracyl phosphoribosyl transferase (*UPRT*) locus. Loss of *UPRT* results in resistance to 5-fluorodeoxyuridine (FUDR), thus allowing the selection of the transgenic parasite.

The cKD HA-TgABCE1 cell line was complemented by adding a copy of the *TgABCE1* gene under the control of a tubulin promoter at the *UPRT* locus. The full-length *TgABCE1* cDNA sequence (1846 bp) was amplified by PCR using primers ML6630 and ML6631 from the plasmid pGBKT7-myc-TgABCE1 to express an N-terminal myc-tagged *TgABCE1* copy. This sequence was then cloned downstream of the tubulin promoter in the pUPRT-TUB vector^72^, resulting in the pUPRT-myc-TgABCE1 plasmid. The plasmid was subsequently linearized with KpnI and BamHI before transfecting the cKD HA-TgABCE1 mutant cell line, together with a plasmid expressing Cas9 and guide RNAs (gRNA), specific to the 5’ and 3’ UTR of the *UPRT l*ocus, under the control of a U6 promoter^73^. The pU6-Cas9-sgRNA plasmids were constructed by hybridizing and cloning the sgRNA primers into the pU6-Cas9 plasmid using BsaI restriction sites. Transgenic parasites were selected using 5 μM 5-Fluoro-2 - deoxyuridine (FUDR, F0503, Sigma-Aldrich) and cloned by serial limiting dilution to obtain the cKD HA-TgABCE1-WT complemented cell line. A similar approach was used to express a myc-tagged TgABCE1 in the context of the TGGT1_218950-mAID-HA_3_ cell line.

For the complementation of the cKD HA-TgABCE1 cell line with the copy of *TgABCE1* containing a mutated cysteine 36, we performed PCR-directed mutagenesis with the Quikchange mutagenesis kit (200515, Agilent), using primers ML6618/ML6619 with the pUPRT-myc-TgABCE1 plasmid as a template. In the case of the complementation with the copy of *TgABCE1* lacking the N-terminal end, the DNA sequence was amplified by using primers ML6848/ML6849 from the vector pUPRT-myc-TgABCE1 to remove the specific N-terminal region. All the constructs were verified by sequencing. Cloning and transfection were performed as previously described for the cKD HA-TgABCE1 WT-complemented cell line. For both cell lines, integration at the *UPRT* locus was verified by PCR using primers ML1428 and ML6765 specific to *TgABCE1*.

### Generating an Auxin-inducible TGGT1_218950 knockdown cell line

The 3’ coding sequence of *TGGT1_218950* was fused to the mAID sequence, followed by sequence coding for a triple HA tag by homology-directed repair using CRISPR/Cas9, resulting of cKD TGGT1_218950-mAID-HA_3_ cell line. The TIR1 ΔKu80 strain was co-transfected with 50 µg of the pU6 -Cas9 carrying the siRNA targeting the 3′ untranslated region (3′UTR) of the *TGGT1_218950* gene and with 20 µg of a PCR product containing the corresponding homology regions and the desired tagging cassette: miniAID sequence and the HA tag followed by the *HXGPRT* cassette was amplified. The mAID-HA-HXGPRT cassette was amplified from the pTUB8YFP-mAID-HA3 plasmid^52^ with the primers ML6905 and ML6906. The pU6-sgRNA plasmids were constructed by hybridizing and cloning the sgRNA primers, ML6895 and ML6896, into the pU6-Cas9 plasmid using BsaI restriction sites. Following transfection, positive transformants were selected for antibiotic resistance by targeting the hypoxanthine-xanthine-guanine phosphoribosyltransferase (HXGPRT) resistance cassette. Clones were isolated by limiting dilution and verified by PCR with the primers ML6903 and ML1476.

### Electron microscopy

Electron microscopy was performed as described earlier^27^. HFFs were infected by tachyzoites at a multiplicity of infection of 1 to 2 parasites per host cell (corresponding to ∼ 2 to 2.10^5^ parasites) in Lab-Tek chamber slides (177437, Nunc). cKD HA-TgABCE1 parasites were pre-cultured with ATc for 24 h, then scraped and allowed to reinvade HFFs in the Lab-Tek slides for an additional 24 h in the presence of ATc or not. Alternatively, RH TATi ΔKu80 parasites were grown on HFFs seeded on Lab-Tek chamber slides, and cycloheximide was used at 100 µg/ml for 6 h. Slides were then chemically fixed for 2 h at room temperature using a fixation solution (2.5% glutaraldehyde in 0.1 M cacodylate buffer, pH 7.4). The samples were stored at 4°C until further processing. Embedding in resin was carried out using a Pelco Biowave PRO+ Microwave processing system (Ted Pella). The details of the program are listed in **Table S4**. Briefly, the cells were first fixed with a double osmium staining procedure as follows: 1) 1% OsO_4_, 5 mM CaCl_2_, in cacodylate buffer 0,1 M pH 7.4; 2) cacodylate buffer washes; 3) In situ reduction with potassium ferricyanide 1.5 %, 5 mM CaCl_2_ in cacodylate buffer; 4) distilled water washes; 5) 1% thiocarbohydrazide (TCH) in water 6) water washes 7) 1% OsO_4_ in water; 8) distilled water washes. The samples were then incubated in 2% uranyl acetate overnight at 4°C. The next day, the samples in uranyl acetate were heated 10 min at 37°C and processed with the Biowave (without washing). After processing, the samples were washed with water, incubated in lead aspartate, washed with water, and then dehydrated in growing concentrations of acetonitrile. Finally, they were progressively impregnated in EPON Hard+ resin directly into the Labtek. The resin was then polymerized at 60°C in an oven for 48 hours. TCH and lead aspartate were prepared extemporaneously as described in Deerinck et al^74^.

Dehydration was performed by increasing the acetonitrile concentration. All chemicals were obtained from Electron Microscopy Sciences. Thin serial sections were prepared using a UCT ultramicrotome (Leica) equipped with an ultra 35° diamond knife (Diatome). Section ribbons were collected on silicon wafers (Ted Pella) and imaged with a Zeiss Gemini 360 scanning electron microscope on the Montpellier ressources imagerie (MRI) EM4Bio facility under high vacuum at 1.5 kV. Final images were acquired using a Sense BSD detector (Zeiss) at a working distance between 3.5 and 4 mm. Single images or mosaics were acquired using Atlas 5 software (Zeiss) with a 6.4 µs dwell time. Pixel size is indicated in the figure legends.

### Immunofluorescence assays (IFA)

For IFAs, coverslips seeded with HFFs were infected with tachyzoites and the host cell monolayer was then fixed with 4% PFA diluted in phosphate-buffered saline (PBS) for 20 min. Cells were washed three times with PBS, permeabilized with 0.3% Triton X-100 in PBS for 10 min, and then saturated with blocking solution (2% Bovine Serum Albumin (BSA) in PBS) for 1 h before immunolabeling with primary antibody for 1 h. After 3 washes in PBS, the corresponding secondary antibody was incubated for 1 h. Coverslips were finally incubated with 1 μg/ml 4,6-diamidino-2-phenylindole (DAPI) for 5 min before 3 washes in PBS and, lastly, mounted using Immu-Mount (9990402, Epredia) onto microscope slides.

Primary antibodies used were prepared in 2% BSA/PBS and used at the following concentrations: rat monoclonal anti-HA (1:1,000, 3F10 Roche), mouse monoclonal anti-myc (1:100, 9E10 Sigma-Aldrich), rabbit anti-IMC3^75^ (1:1,000), mouse anti-SAG1^76^ (T41E5, 1:1,000), and mouse anti-puromycin (1:1,000, 12D10, MABE943 Sigma-Aldrich). Highly cross-adsorbed Alexa Fluor 488-conjugated or Alexa Fluor 594-conjugated anti-rat-, rabbit-, or mouse-IgG secondary antibodies were all from ThermoFisher and were diluted at 1:4,000. Cyst wall was stained with biotin-labeled *Dolichos Biflorus* lectin (1:300, L-6533, Sigma-Aldrich) and detected with FITC-conjugated streptavidin (1:300, Invitrogen,). LDs were stained with Nile Red (1 μg/ml, SNN1008, Sigma-Aldrich) or with BODIPY 493/503 (10 µM, HY-W090090, MedChemExpress) for 30 min.

For monitoring stage conversion, *Dolichos biflorus* lectin (DBL) was used for the detection of cyst walls as described previously and the ME49 type II cell line was used as a control upon pH-driven conversion^77^.

All images were acquired with a Zeiss Axio Observer epifluorescence inverted microscope equipped with a Zeiss Axiocam 712 camera. Images were processed on Zen v3.6 (Blue edition) software (Zeiss). Z-stack acquisitions were processed by maximum intensity orthogonal projection when assessing LD number and area. Adjustments of brightness and contrast were applied uniformly and paired images were acquired with the same exposure time.

### Host cell lipid uptake assay

HFF cells were pretreated with oleic acid (HY-N1446, MedChemExpress) at 400 µM and BODIPY 493/503 at 10 µM for 18h, washed five times with PBS and then incubated in complete DMEM medium for 30 min. Then, the cells were incubated with oleic acid at 200 µM and infected with cKD HA-ABCE1 tachyzoites either in the presence or absence of ATc for 48 h or with microtubule-disrupting agent oryzalin (HY-147092, MedChemExpress) at 2.5 µM for 48 h.

Upon treatment with 2.5 µM of oryzalin for 48 h, the compound was removed by washes in HBSS and the parasites were then maintained in complete medium for 48 h. The parasites were fixed and permeabilized according to the IFA protocol, after which the LDs were stained with BODIPY493/503 for 30 min.

### Plaque and replication assays

Tachyzoites from the cKD HA-TgABCE1, the cKD TGGT1_218950-mAID-HA_3_ transgenic cell lines or parental strain (TATi ΔKu80) were allowed to invade monolayers of HFFs in the presence or absence of ATc or IAA. Parasites were cultivated for 7 days at 37°C and 5% CO_2_ and fixed with 4% (w/v) PFA (diluted in PBS) for 20 min. Cells were stained with 0.1% crystal violet solution (V5265, Sigma-Aldrich), washed, and air-dried before imaging on an Olympus MVX10 microscope. For the reversibility assay, drug washout was carefully performed after 7 days of ATc pretreatment and cultures were kept for another 7 days of growth, non-treated control conditions were also infected at this time point. For the viability assay, parasites were pre-treated in a flask with ATc for 48 h. The parasites were then incubated on ice for 20 min, followed by a 20 min invasion period at 37°C, after which the medium was washed and the culture was maintained for 7 days.

For the replication assay, HFF monolayers grown on coverslips were infected with 2 × 10^5^ freshly egressed parasites, pre-treated or not with ATc for 24 h (or IAA for 24 h). Invasion was allowed for 2 h, HFFs were then washed 3 times with HBSS to remove extracellular parasites incapable of entering host cells, and intracellular parasites were left to replicate for 24 h at 37°C in the presence or not of ATc or IAA. Cells were then fixed, permeabilized, and parasites were immunodetected by IFA with mouse anti-SAG1^76^ antibody. The number of parasites per vacuole was scored. Independent experiments were conducted 3 times, and 100 random vacuoles were counted for each condition.

### DNA content analysis

DNA content was analyzed by flow cytometry as described previously^39^. Parasites from the cKD HA-TgABCE1 and TATi ΔKu80 cell lines were grown in the presence or absence of ATc and intracellular parasites were released from host cells by scraping, then passage through 26G needles and finally passed on a 40 µm filter and fixed overnight in a solution of 70% ethanol and 30% PBS at 4°C. After fixation, parasites were washed in PBS and stained with a 30 µM propidium iodide solution for 30 min. DNA content was then quantified by flow cytometry with an Aurora cytometer (Cytek) from the MRI facility and analyzed with FlowJo v10 (https://www.flowjo.com/, BD Biosciences). Raw FCS files and gating strategy for these cytometry experiments are available at https://figshare.com/s/7fc652dc3cc7d541942b.

### Immunoblotting and antibodies

Protein extracts were prepared from 10^7^ extracellular parasites resuspended in SDS-PAGE sample loading buffer supplemented with 100 mM dithiothreitol (Sigma-Aldrich) at a final concentration of 10^6^ parasites/μl. Extracts were treated with benzonase (E1014 Sigma-Aldrich) to remove DNA from samples. They were incubated for 30 min at 37°C and then boiled at 95°C for 10 min. Proteins were separated on a 10% SDS-PAGE gel (10% acrylamide/bis, 0.4 M Tris–HCl pH 8.8, 0.1% SDS, and 0.1% APS, TEMED) before being transferred on 0.45 µm nitrocellulose membrane (GE10600002, Amersham Protran, GE Healthcare Life Science) for subsequent protein detection. Blots were blocked with 5% dried milk in TNT buffer (0.1 M Tris-HCl (pH 7.6), 0.15 M NaCl, 0.05%, Tween 20).

Primary antibodies used for immunodetection were resuspended at their respective working concentration in 5% milk/TNT buffer. Antibodies used in this study for immunoblot detection were rat anti-HA (1:1,000, Roche), mouse anti-myc (1:100, 9E10 Sigma-Aldrich), mouse anti-SAG1^76^ (1:50, hybridoma), mouse anti-actin^78^ (1:25, hybridoma) and mouse anti-puromycin (1:1,000, 12D10, MABE943 Sigma-Aldrich). Alkaline phosphatase (AP)-conjugated anti-mouse-, or anti rabbit-IgG were from Promega and used at 1:7,500. AP-conjugated anti-rat-IgG secondary antibodies were from Sigma-Aldrich and used at 1:10 000. Proteins were visualized with BCIP/NBT development substrates (Promega). Alternatively, horseradish peroxidase (HRP)-conjugated anti rat- or mouse-IgG secondary antibodies (all from ThermoFisher, diluted at 1:10,000) were used and signal was revealed with the Clarity Max™ Western ECL (Bio-Rad) substrates.

### Puromycin labeling

Cell lines were cultured for 2 days in the presence or absence of ATc or up to 24 h with the auxin indole-3-acetic acid (Sigma-Aldrich). Freshly egressed parasites were filtered on 40 μm Cell Strainer (723–2757, VWR). After filtration and counting, parasites were treated with puromycin (100 μg/ml, puromycin dihydrochloride, Sigma-Aldrich) for 15 min at 37°C and 5% CO_2_. For translation inhibition control, parasites were treated with cycloheximide (100 μg/ml, Sigma-Aldrich) for 10 min before puromycin incubation. After treatment, parasites were washed in DPBS (Dulbecco’s phosphate-buffered saline, Gibco) and collected by centrifugation. The pellet was resuspended in Laemmli buffer and separated on Mini-Protean TGX Stain-free gels 10% (BioRad) activated by a UV-induced 1-min reaction to produce tryptophan residue fluorescence to allow for global protein quantification, following manufacturer instructions. Proteins were then transferred to a nitrocellulose membrane for immunodetection using mouse anti-puromycin (1:1,000, 12D10, MABE943, Sigma-Aldrich). Both signals were quantified by densitometry using the ImageJ software.

### Parasite viability assay

Trypan blue viability assays were conducted as described previously^36^. Briefly, freshly egressed extracellular parasites (previously cultivated in the presence or absence of ATc for 48 h, or IAA for 24 h) were incubated in complete medium at 37 °C with 5% CO_2_ for 6 h and stained with Trypan Blue (TB, Sigma-Aldrich) to determine the mortality rate by counting TB-positive parasites under a fluorescence microscope (excitation: 550/25 nm, emission: 605/70 nm). Typically, dead parasites take up TB into the cytoplasm because of loss of their membrane integrity, and there TB becomes fluorescent as it is complexed to proteins^79^.

### Protein labeling with L-azidohomoalanine and mass spectrometry quantification

To label newly synthesized proteins, we used the L-azidohomoalanine (AHA) methionine analog (C10102, Invitrogen), which can be chemoselectively linked to an alkyne-conjugated moiety via copper-catalyzed azide-alkyne cycloaddition reaction (CuAAC click reaction). Before starting the labeling, the cKD HA-TgABCE1 intracellular parasites, pre-treated or not with ATc for 48 h, were pre-incubated in methionine-free DMEM (Gibco, 21013024). Then, the cells were labeled for 6 h at 37°C and 5% CO2 with 25 µM AHA (Click-iT AHA, C10102 Invitrogen) in methionine-free DMEM (21013024, Gibco) as described before^36^, to allow for the incorporation into newly synthesized proteins in each sample. Protein synthesis inhibitor cycloheximide (100 µg/ml, Sigma-Aldrich) was used in parallel for 6 h as a negative control. To confirm the incorporation of the AHA label into newly synthesized proteins, we performed a CuAAC click reaction with Alexa-Fluor-488-azide directly on fixed coverslips, followed by fluorescence microscopy imaging. The treated cells were washed with PBS, fixed with 4% paraformaldehyde-PBS for 15 min, and subsequently permeabilized with 0.3% Triton X-100 in PBS for 10 to 15 min. For the CuAAC click reaction, the coverslips were incubated for 60 min at room temperature in the dark with a reaction cocktail containing Alexa Fluor 488 azide (A10266, Invitrogen), 1 mM of CuSO₄,,100 µM tris(benzyltriazolylmethyl)amine (TBTA) diluted in DMSO and 1 mM of tris(2 carboxyethyl)phosphine (TCEP) diluted in H_2_0. After three washes with PBS, DNA was counterstained with DAPI, and the slides were mounted for fluorescence microscopy. For mass spectrometry analysis, after treatment, intracellular parasites were released by scraping of the host cells and 3 passages through a 26G needle. To eliminate cell host debris, parasites were filtered through 40µm Cell Strainer (723–2757, VWR) and harvested by centrifugation at 650 g for 5 min. The pellets were then resuspended in Urea Lysis Buffer (C10416, Invitrogen) supplemented with cOmplete Mini protease inhibitors mix (11836170001, Roche) and sonicated for 3 times for 30 seconds at 70% of amplitude in ice. Then, the click reaction was performed directly on alkyne agarose resin according to the manufacturer’s protocol for the Click-iT protein Enrichment kit (C10416, Invitrogen) for click chemistry capture of azide-modified proteins. AHA-labeled proteins were then digested with trypsin and analyzed by mass spectrometry.

Briefly, samples mixed with beads were dispersed in 200 µl of 100 mM Tris / 2 mM NaCl₂ / 10% ACN buffer. Then, 10 µl of trypsin solution at a concentration 0.1 µg/µl (Sequencing Grade Modified Trypsin, Promega) was added. The mixture was vortexed for 5 min., and 50 µl of buffer was added along the tube walls to bring down the beads sticking to the sides. Digestion was carried out overnight at 37°C. The samples were then centrifuged for 5 min. at 1000g, and the supernatants were transferred to a 2 ml Eppendorf tube. 500 µl of LC-MS/MS-grade water was added to the beads, vortexed for 5 min., and centrifuged again for 5 min. at 1000 g. The resulting supernatant was pooled with the previous fraction. Finally, 1 ml of water was added to each sample, acidified with 2 µl of pure TFA, and centrifuged for an additional 5 min. at 1000 g. Then, peptides were purified using a C18 cartridge (Sep-Pak R Vac 1cc, WAT036820, Waters Corporation. The cartridge was conditioned with 1 ml of 50% ACN / 0.1% TFA, followed by two washes with 1 ml of 0.1% TFA. Supernatants with peptides were loaded onto the cartridge. They were then washed twice with 1 ml of 0.1% TFA and finally eluted with 700 µl of 50% ACN / 0.1% TFA. The eluate was dried in a vacuum centrifuge until complete evaporation. Peptides were resuspended in 6 µl of 2% FA before LC-MS/MS analysis.

The LC-MS/MS experiments were performed using an Ultimate 3000 RSLC nano system (ThermoFisher Scientific) interfaced online with a nano easy ion source and a Exploris 240 Orbitrap mass spectrometer (ThermoFisher Scientific). The samples were analyzed in Data Dependent Acquisition (DDA). 6µl of peptide solution was first loaded onto a pre-column (Thermo Scientific PepMap 100 C18, 5 µm particle size, 100 Å pore size, 300 µm i.d. x 5 mm length) from the Ultimate 3000 autosampler with 0.05% TFA in water at a flow rate of 10 µL/min. The peptides were separated by reverse-phase column (Thermo Scientific PepMap C18, 2 *μ*m particle size, 100 Å pore size, 75 *μ*m i.d. 50 cm length) at a flow rate of 300 nl/min. After a 3 min loading period, the column valve was switched to allow elution of peptides from the pre-column onto the analytical column. Loading buffer (solvent A) was 0.1% FA in water, and elution buffer (solvent B) was 0.1% FA in 80% acetonitrile. The linear gradient employed was 2-25% of solvent B in 103 min, then 25-40% of solvent B from 103 to 123 min., finally 40-90% of solvent B from 123 to 125 min. The total run time was 150 min. including a high organic wash step and re-equilibration step. Peptides were transferred to the gaseous phase with positive ion electrospray ionisation at 1.9 kV. In DDA the top 15 precursors were acquired between 350 and 1200 m/z with a 2 Th (Thomson) selection window, dynamic exclusion of 40 s, normalized collision energy (NCE) of 30, and resolutions of 120 000 for MS and 15 000 for MS2. Spectra were recorded with Xcalibur software (4.7.69.37) (ThermoFisher Scientific).

The .raw files were analysed with MaxQuant version 2.0.3 using default settings. The minimal peptide length was set to 6. The criteria “Trypsin/P” (which means C-terminus peptides of “K/R” unless followed by “P”: “K/R” followed by “P” cannot be a cleavage site) was chosen as the digestion enzyme. Carbamidomethylation of cysteine was selected as a fixed modification and as variable modification: N-terminal acetylation of protein; oxidation of methionine; N-terminal-pyroglutamylation of glutamine and glutamate, and substitution of Methionine with AHA. Up to two missed cleavages were allowed. The mass tolerance for the precursor was 20 and 4.5 ppm for the first and the main searches, respectively, and for the fragment ions it was 20 ppm.

The files were searched against the *T. gondii* proteome (March 2020 -https://www.uniprot.org/proteomes/UP000005641-8450 entries) and the Maxquant contaminant database. Identified proteins were filtered according to the following criteria: at least two different trypsin peptides with at least one unique peptide, an E value below 0.01 and a protein E value smaller than 0.01 were required. Using the above criteria, the rate of false peptide sequence assignment and false protein identification were lower than 1%.

### Co-immunoprecipitation and mass spectrometry identification

Parasites of the cKD HA-TgABCE1 transgenic cell and the parental strain line TATi ΔKu80 were treated for 2 days. After treatment, intracellular parasites were released by scraping of the host cells and 3 passages through a 26G needle. To eliminate cell host debris, parasites were filtered through 40 μm Cell Strainer (723–2757, VWR) and harvested by centrifugation at 650 g for 5 min. Parasites were resuspended in cold lysis buffer (1%NP40, 50 mM Tris-HCl (pH 8), 150 mM NaCl, 4 mM EDTA, supplemented with cOmplete Mini protease inhibitors mix (11836170001, Roche) and incubated overnight at 4°C on a rotating wheel. Centrifugation of insoluble material was performed at 13,500 g for 30 min at 4°C. The supernatant was transferred to a tube containing 40 μl of anti-HA magnetic beads (88836, ThermoFisher) for 4 h at 4°C on a rotating wheel. The depleted fraction was then removed and beads were washed 5 times with cold lysis buffer. For the elution of immunoprecipitated proteins, the beads were directly eluted with 30 µL of Laemmli buffer at 95°C for 5 min and resolved on a 10% SDS-PAGE for 35 min at 100V. In-gel proteins were stained with PageBlue Protein Staining Solution (24620, Thermo Scientific). Each lane was cut into 3 identical pieces, which were processed for mass spectrometry as described previously^21,80^.

All raw MS data and MaxQuant files generated have been deposited to the ProteomeXchange Consortium via the PRIDE partner repository (https://www.ebi.ac.uk/pride/archive) with the dataset identifiers PXD080321 and PXD080322.

### Bioinformatic analyses

Sequence alignments were performed using the CLUSTALW algorithm of the Geneious 6.1.8 software suite (http://www.geneious.com). Motif analysis was performed with the Multiple Em for Motif Elicitation (MEME) suite (https://meme-suite.org/meme/). Nuclear localization signal predictions were performed using NLSmapper (https://nls-mapper.iab.keio.ac.jp/cgi-bin/NLS_Mapper_y.cgi) and NLSexplorer (http://www.csbio.sjtu.edu.cn/bioinf/NLSExplorer/). The HHpred server was used to detect remote structural homology (https://toolkit.tuebingen.mpg.de/tools/hhpred). The AlphaFold^81^ server (https://alphafold.ebi.ac.uk/) was used for structural prediction, and structures were rendered using ChimeraX v 1.11^82^.

### Statistical analyses

Statistical analyses were performed with the Prism 8.3 software (Graphpad). Values were expressed as means ± standard deviation (SD) and statistical significance was assessed using Student’s unpaired t-test or ANOVA, as indicated in the figure legends. Sample sizes (*n*) correspond to independent biological replicates unless otherwise stated and all experiments were performed with at least three independent biological replicates. Cell morphology phenotypes were usually scored in a blinded manner. Experiments lacking statistical analysis (like representative immunoblots or microscopy images) were usually repeated at least three times and found to yield similar results.

## Acknowledgements

We thank V. Carruthers, S. Lourido, M.J. Gubbels, J.F. Dubremetz, D. Soldati-Favre, B. Striepen, and D. Sibley for providing cell lines and reagents. Mass spectrometry experiments were carried out using the facilities of the Montpellier Proteomics Platform (PPM, MSPP site BioCampus Montpellier), a member of the national Proteomics French Infrastructure (ProFI UAR 2048) supported by the French National Research Agency (ANR-24-INBS-0015, Investments for the future F2030). We also acknowledge the MRI imaging and cytometry facility, member of the national infrastructure France-BioImaging (https://ror.org/01y7vt929) supported by the French National Research Agency (ANR-24-INBS-0005 FBI BIOGEN). This work was supported by grants from the Agence Nationale de la Recherche (ANR-22-CE20-0026 and ANR-24-CE15-3620) to S.B. M.G.D. is supported by a doctoral fellowship from the Fondation pour la Recherche Médicale.

## Author contributions

A.J.M.M., M.G.D., E.A.R. and S.B. performed most of the experiments and analyzed the data. V.R. and V.D. performed the mass spectrometry analyses. A.G. and L.B. performed the electron microscopy experiments. S.B. conceived and supervised the study and acquired funding. S.B. wrote the first draft of the manuscript with contribution of A.J.M.M. to subsequent versions and revisions. All authors reviewed, commented on, and approved the manuscript.

## Supplementary table legends

**Table S1. Peptide quantification by mass spectrometry upon TgABCE1 depletion (ATc condition) or global protein synthesis inhibition (cycloheximide, CHX condition).** Proteins were enriched by click-chemistry after L-azidohomoalanine incorporation. Data displayed are from *n* = 3 independent biological replicates.

**Table S2. Proteins identified as co-immunoprecipitating with TgABCE1 through comparative mass spectrometry analysis of immunoprecipitated extracts.** Data are from *n* = 3 independent biological replicates.

**Table S3. Primers used in this study.**

**Table 4. Microwave processing program for the preparation of electron microscopy samples.**

