## Supplemental Figures S1 to S9 for "ABCE1-dependent translational control links Fe-S cluster biogenesis to parasite growth and lipid homeostasis in *Toxoplasma gondii*"

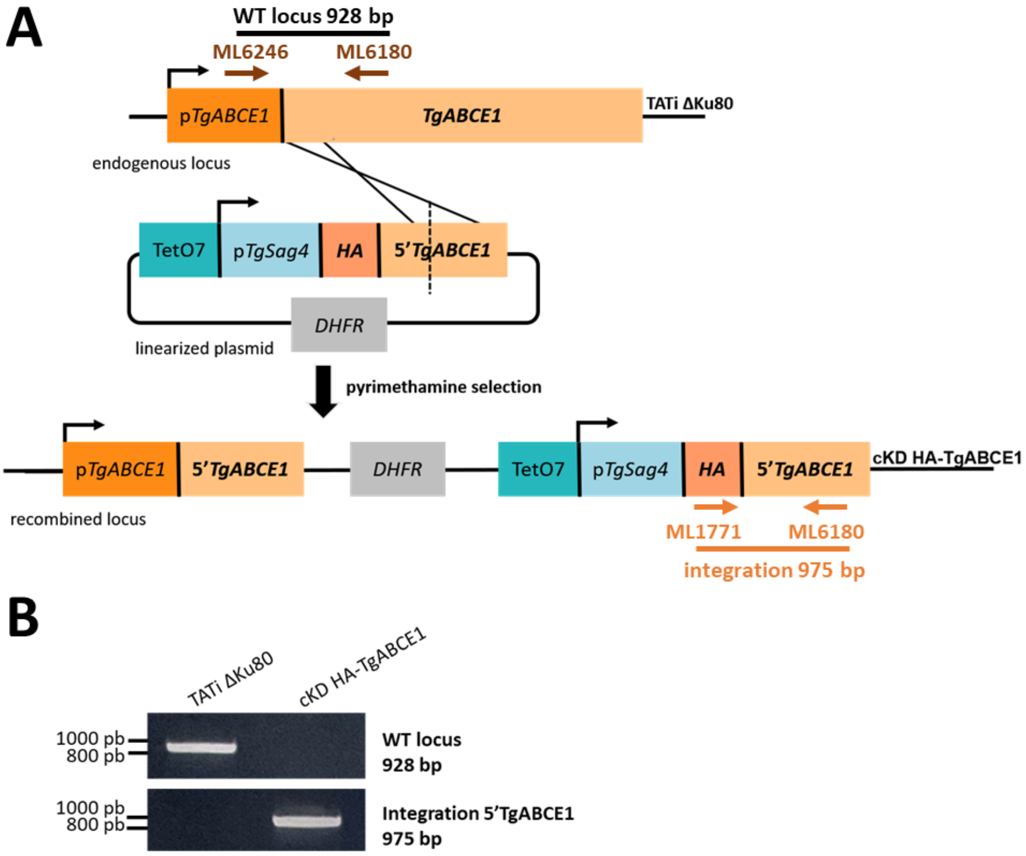


**Figure S1. Generation of the cKD HA-TgABCE1 cell line. A.** Strategy for generating the inducible knockdown of *TgABCE1* by promoter replacement and simultaneous N-terminal tagging of the TgABCE1 protein in the TATi ΔKu80 cell line. The endogenous *TgABCE1* promoter was replaced with an ATc-regulatable TetOSag4 promoter by homologous recombination. Positive selection with pyrimethamine was used to select cKD HA-TgABCE1 transgenic parasites based on the DHFR resistance cassette. **B.** Diagnostic PCR for either detecting the WT *TgABCE1* locus or the recombined 5' with primers labelled on **A**.

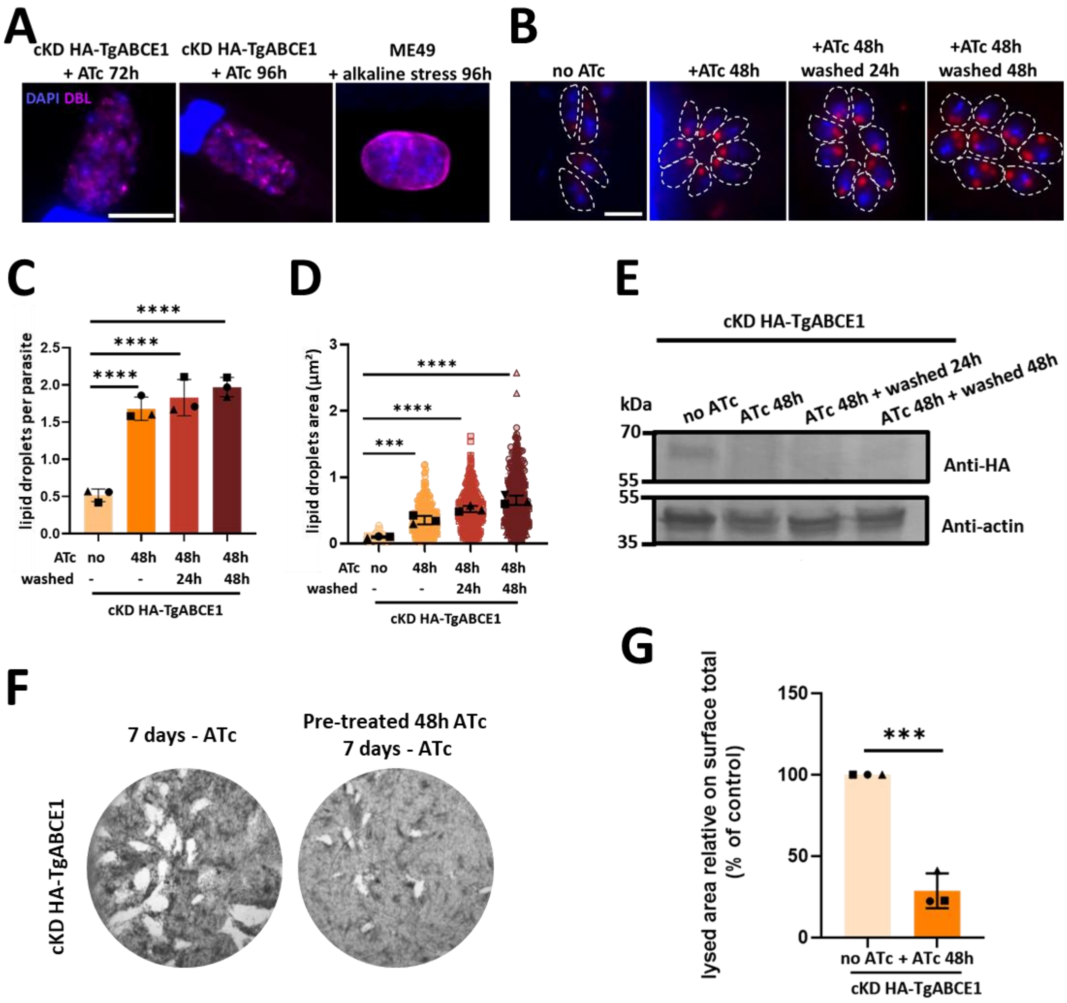


**Figure S2. TgABCE1 depletion impacts parasite fitness but does not trigger stage conversion. A.** *Dolichos biflorus* lectin (DBL) was used to probe for cyst wall formation upon TgABCE1 depletion for up to 96 h and showed no peripheral labeling in contrast to the ME49 positive control left to differentiate following alkaline stress for 96 h. DNA is stained with DAPI. Scale bar: 10 µm. **B.** Nile red staining showed that lipid droplets (LDs) accumulation is irreversible in the cKD HA-TgABCE1 mutant. The parasites were treated with ATc for 48 h, then the drug was washed out with PBS, and the parasites were cultured in complete DMEM medium without ATc for 24 or 48 h. Parasite shapes are outlined with dashed lines, DNA is stained with DAPI. Scale bar: 5 µm. **C, D.** Quantification of the number and surface area of LDs as observed in **A.**, respectively; 100 parasites were analyzed per condition. Values are presented as mean ± standard deviation of $n = 3$ independent biological replicates (different symbols represent different series): *** $p$-value ≤ 0.001; **** $p$-value ≤ 0.0001. $p$-values derived from a one-way ANOVA with Dunnett's multiple comparison test. **E.** Western blot showing that TgABCE1 depletion is irreversible after 48 h of depletion with ATc. cKD HA-TgABCE1 parasites were cultured in the presence or absence of ATc for 48 h, after which the ATc was removed through repeated washing and then parasites were grown further for 24 or 48 h. Actin was used as a loading control. **F.** Plaque assay was performed by infecting a monolayer of HFF cells with cKD HA-TgABCE1 cells pre-treated with ATc for 48 h. The parasites invaded the monolayer for 20 min. at 37°C. After washing with PBS, the cell lines were cultured for 7 days in the absence of ATc. **G.** Quantification of the plaque area observed in **F.** Results are expressed as the percentage of lysed area relative to the total lysed surface of control (cKD HA-TgABCE1, set as 100% for reference). Values represented are mean ± standard deviation from $n$ = 3 independent biological replicates, *** $p$-value ≤0.001, Student's t-test.

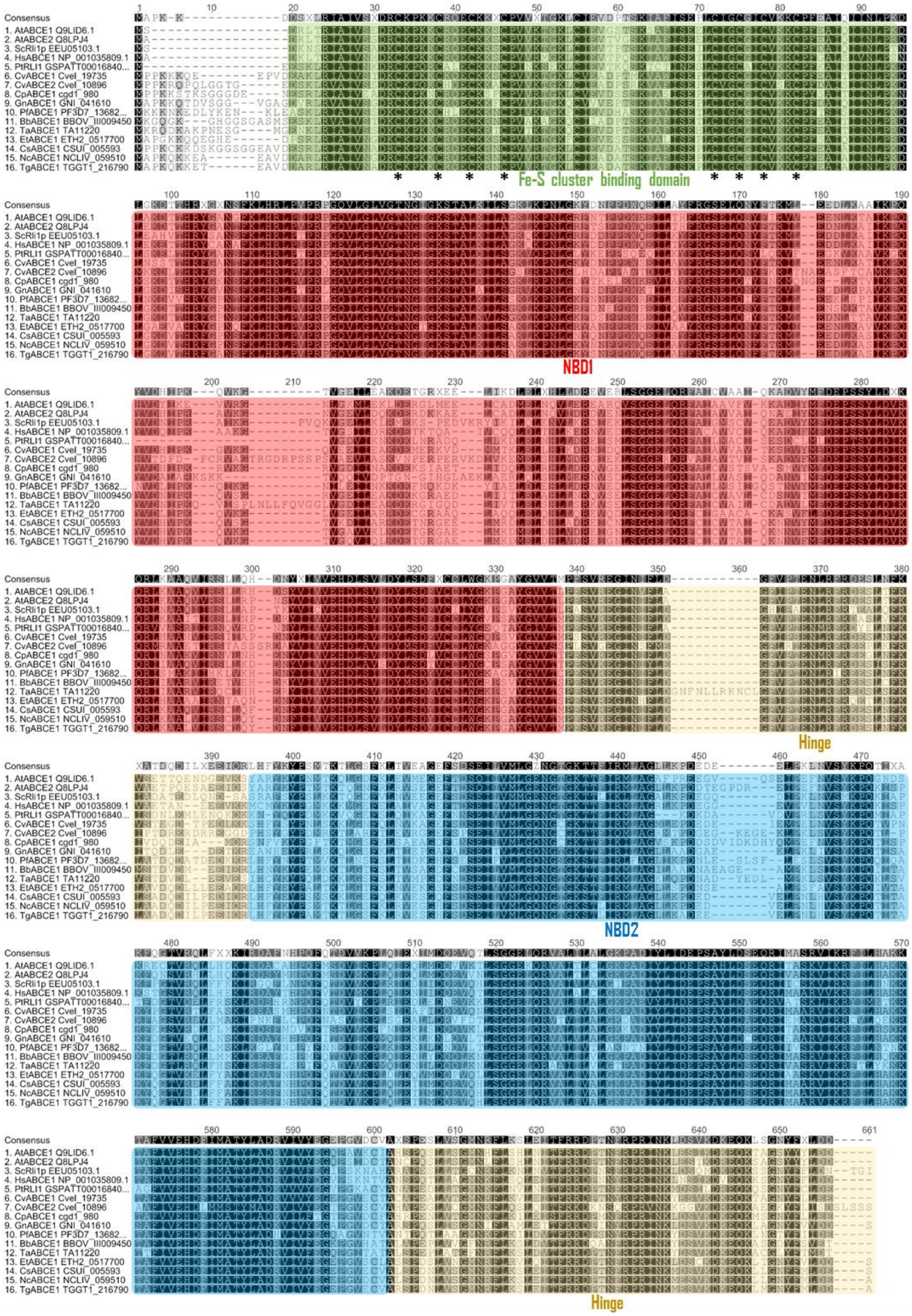


**Figure S3. CLUSTALW alignment of the primary protein sequence for ABCE1/ABCE2 homologs of various eukaryotes**

GenBank or VeupathDB accession numbers are listed for each protein. Different domains are highlighted in color : Fe-S cluster binding region (green), nucleotide binding domains 1 (NBD1, red) and 2 (NBD2, blue), as well as hinge regions (yellow). Asterisks mark cysteines potentially involved in coordinating Fe-S clusters.

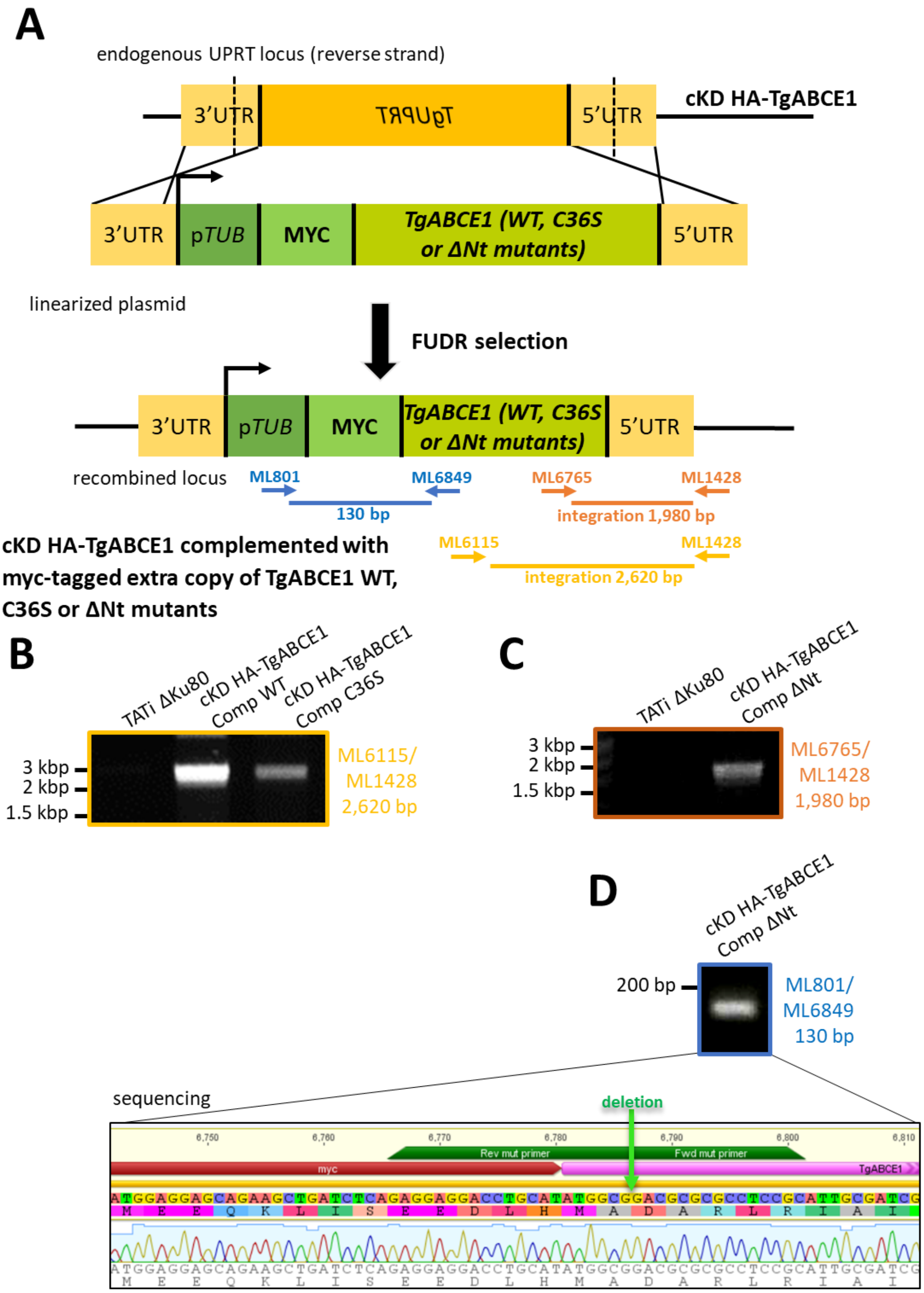


**Figure S4. Generating constructs for TgABCE1 complementation**

**A.** Schematic representation of the strategy for generating cell lines expressing myc-tagged wild-type (WT), cysteine 36 mutation to serine (C36S), or N-terminally truncated (ΔNter) copies of TgABCE1 by integrating the sequence coding for the extra copy of the gene of interest by double homologous recombination at the *Uracil Phosphoribosyltransferase* (*UPRT*) locus. Negative selection with 5-fluorodeoxyuridine

(FUDR) was used to select transgenic parasites based on their absence of UPRT expression. **B.** Diagnostic PCR for verifying integration at the *UPRT* locus thanks to the primers described in **A** for the WT and C36S complementations. **C.** Diagnostic PCR for verifying integration at the *UPRT* locus thanks to the primers described in **A** for the ΔNter complementation. **D.** Verification of the N-terminal sequence truncation by amplification of the region by PCR and sequencing of the PCR fragment.

**A**

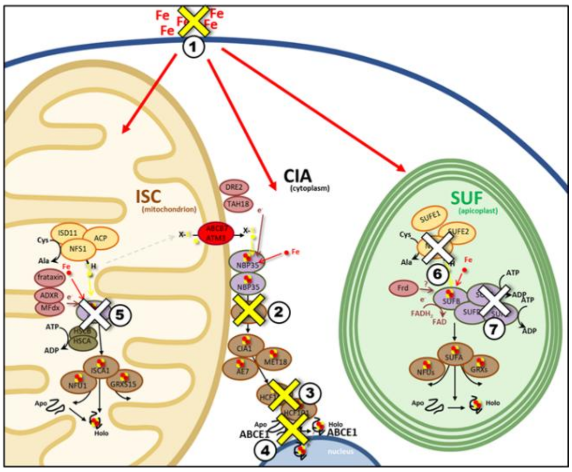


**B**

| | LD number | LD area | reference |
|---|---|---|---|
| RH control | 0.38 ± 0.04 | 0.19 ± 0.04 | Renaud et al 2024 |
| **1.** RH + Fe chelator | 1.06 ± 0.07 | 0.46 ± 0.04 | Renaud et al 2024 |
| **2.** TgNAR1-depleted | 1.56 ± 0.11 | 0.22 ± 0.01 | Renaud et al 2025 |
| **3.** TgHCF101-depleted | 1.24 ± 0.17 | 0.38 ± 0.02 | Renaud et al 2025 |
| **4.** TgABCE1-depleted | 1.92 ± 0.17 | 0.42 ± 0.01 | this work |
| **5.** TgISU1-depleted | 0.58 ± 0.12 | 0.18 ± 0.01 | Renaud et al 2025 |
| **6.** TgNFS2-depleted | 0.42 ± 0.10 | 0.20 ± 0.02 | Renaud et al 2025 |
| **7.** TgSUFC-depleted | 0.60 ± 0.06 | 0.17 ± 0.01 | Renaud et al 2025 |

**Figure S5. Iron chelation or disruption of the CIA Fe–S assembly pathway leads to lipid droplet accumulation in *T. gondii* and may converge on TgABCE1 function**

**A.** Schematic representation of the Fe-S assembly pathways of *T. gondii* describing different levels of interference with Fe-S assembly. Some are leading to lipid droplet (LD) accumulation (yellow crosses) contrarily to others (white crosses). **B.** Table summarizing LD number and area for the different steps illustrated and numbered in **A**.

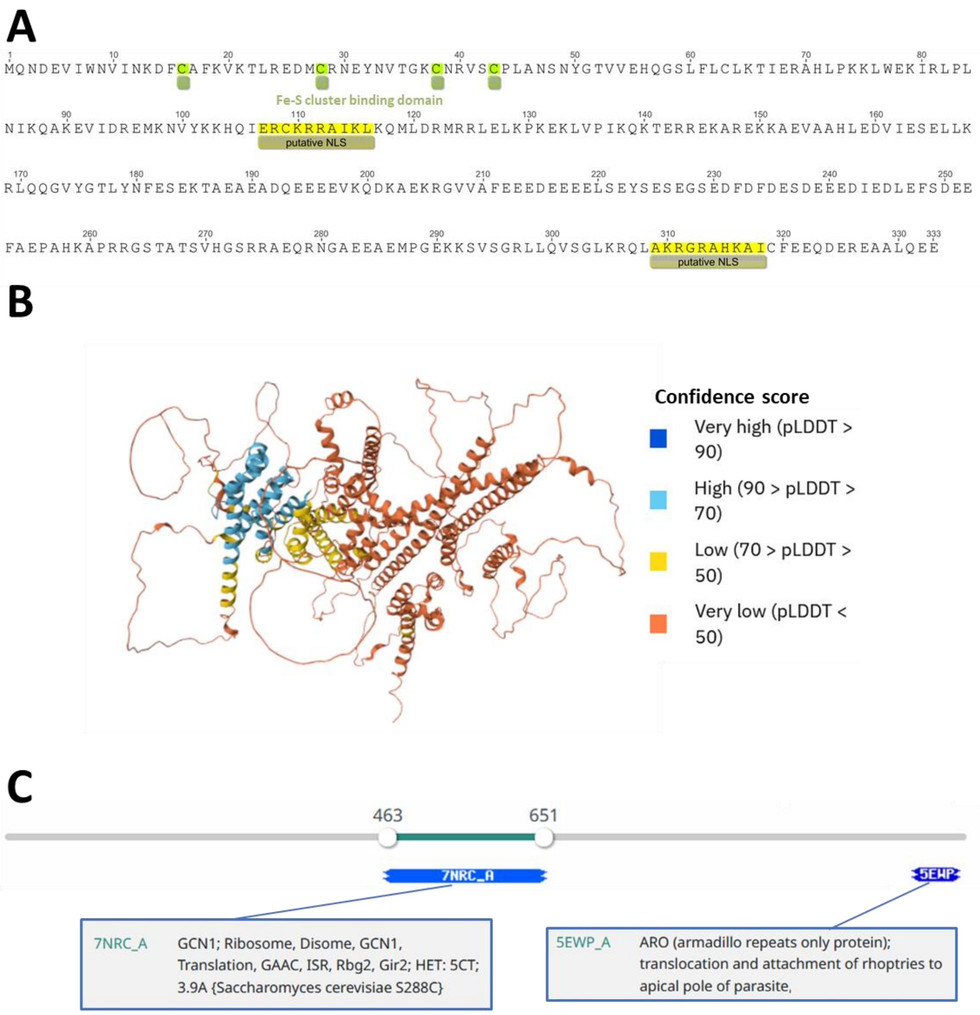


**Figure S6. Sequence and structural information of putative TgABCE1 partners**

**A.** Primary amino acid sequence of TgMAK16, with annotated conserved cysteines potentially involved in Fe-S cluster coordination, and putative nuclear localization signals (NLS). **B.** AlphaFold structural model of TGGT1_218950 showing extensive α-helical repeats. **C.** Motifs with structural homology were detected by analysis with the HHpred server and include homology with GCN1 that is involved in ribosomal quality control.

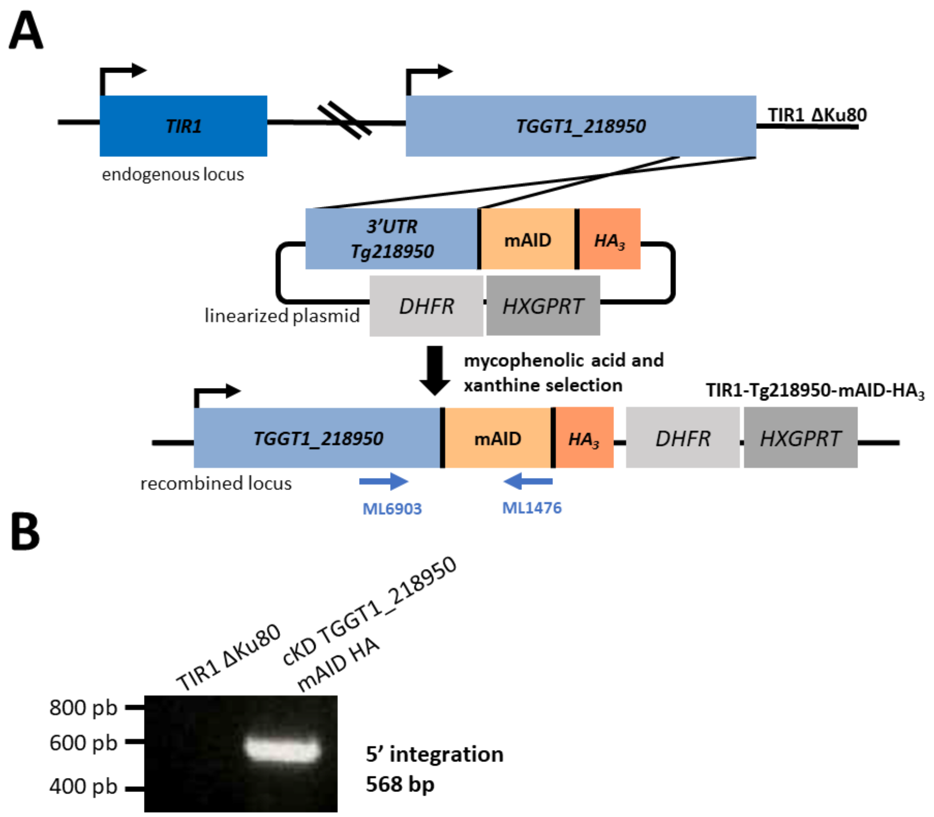


**Figure S7. Generation of a TGGT1_218950 conditional knock down cell line**

**A.** Schematic representation of the strategy for generating the cKD TGGT1_218950-mAID-$HA_3$ cell line by modification of the 5' of the *TGGT1_218950* gene to insert a sequence coding for a triple HA tag and a mini- auxin-inducible degron (mAID) cassette.

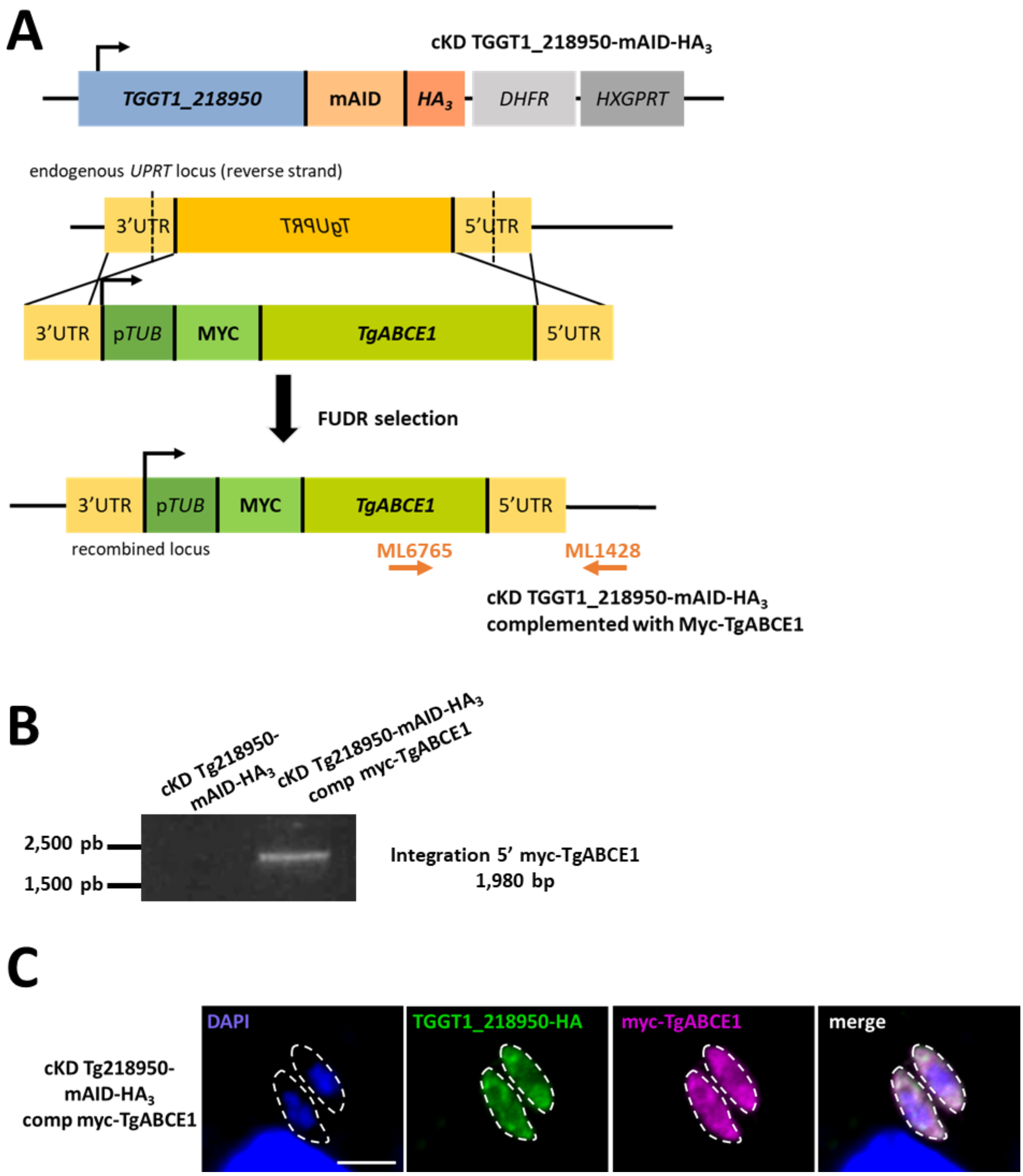


**Figure S8. Generation of a cell line expressing HA-tagged TGGT1_218950 and myc-tagged TgABCE1**

**A.** Schematic representation of the strategy for generating a cell line expressing HA-tagged TGGT1_218950 and myc-tagged TgABCE1. A myc-tagged extra copy of TgABCE1 was expressed at the *Uracil Phosphoribosyltransferase* (*UPRT*) locus in the cKD TGGT1_218950-mAID-$HA_3$ cell line background. **B.** Diagnostic PCR for integration using the primers described in **A. C.** Immunofluorescence with anti-myc antibody confirms the cytoplasmic localization of the extra copies of myc-TgABCE1 (in red) in the cKD Tg218950-maiD-$HA_3$ (in green). Parasites shapes are outlined. DNA was stained with DAPI. Scale bar = 5 µm.

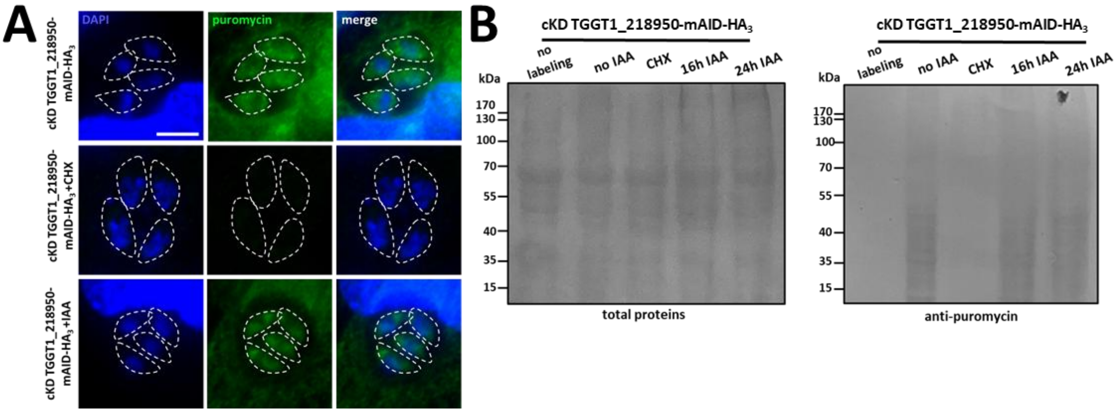


**Figure S9. Depletion of TGGT1_218950 does not strongly impact protein synthesis**

**A.** cKD TGGT1_218950-mAID-HA$_3$ parasites were incubated with cycloheximide (CHX) or indole-3-acetic acid (IAA) for 6h and pulse-labelled with puromycin before being analysed by immunofluorescence microscopy. Parasites shapes are outlined. DNA was stained with DAPI. Scale bar = 5 µm. **B.** Immunoblot analysis of puromycin labeling in the cKD TGGT1_218950-mAID-HA$_3$ parasites showing no strong impact of protein depletion on puromycin signal. Total proteins were stained with Ponceau red.
