## Supplemental Tables S1 to S4 for "ABCE1-dependent translational control links Fe-S cluster biogenesis to parasite growth and lipid homeostasis in *Toxoplasma gondii*": Table S4.docx

**Ferri = K Ferrocyanid; TCH : thiocarbohydrazide; UA = uranyle acetate**

| **Day 1** | | | | | | | | | | | | |  |
| --- | --- | --- | --- | --- | --- | --- | --- | --- | --- | --- | --- | --- | --- |
| **Step name** | | | **Step number** | | **Time** | | **Watts** | **Vent time**  **(sec)** | **Vac time**  **(sec)** | **Set vac**  **(inch of Mercury)** | **load**  **cooler T(°C)** | **user**  **prompt** |  |
| Buffer Rinse | | | 1 | | 40 sec | | 250 | 0 | 0 | 0 | 30 | yes |  |
| Buffer Rinse | | | 2 | | 40 sec | | 250 | 0 | 0 | 0 | 30 | yes |  |
| Buffer Rinse | | | 3 | | 40 sec | | 250 | 0 | 0 | 0 | 30 | yes |  |
| OsO4 ON | | | 4 | | 2 min | | 100 | 30 | 30 | 20 | 30 | yes |  |
| OsO4 OFF | | | 5 | | 2 min | | 0 | 30 | 30 | 20 | 30 | No |  |
| OsO4 ON | | | 6 | | 2 min | | 100 | 30 | 30 | 20 | 30 | No |  |
| OsO4 OFF | | | 7 | | 2 min | | 0 | 30 | 30 | 20 | 30 | No |  |
| OsO4 ON | | | 8 | | 2 min | | 100 | 30 | 30 | 20 | 30 | No |  |
| OsO4 OFF | | | 9 | | 2 min | | 0 | 30 | 30 | 20 | 30 | No |  |
| OsO4 ON | | | 10 | | 2 min | | 100 | 30 | 30 | 20 | 30 | No |  |
| Buffer Rinse | | | 11 | | 40 sec | | 250 | 0 | 0 | 0 | 30 | yes |  |
| Buffer Rinse | | | 12 | | 40 sec | | 250 | 0 | 0 | 0 | 30 | yes |  |
| Buffer Rinse | | | 13 | | 40 sec | | 250 | 0 | 0 | 0 | 30 | yes |  |
| Ferri ON | | | 14 | | 2 min | | 100 | 30 | 30 | 20 | 30 | yes |  |
| Ferri OFF | | | 15 | | 2 min | | 0 | 30 | 30 | 20 | 30 | No |  |
| Ferri ON | | | 16 | | 2 min | | 100 | 30 | 30 | 20 | 30 | No |  |
| Ferri OFF | | | 17 | | 2 min | | 0 | 30 | 30 | 20 | 30 | No |  |
| Ferri ON | | | 18 | | 2 min | | 100 | 30 | 30 | 20 | 30 | No |  |
| Ferri OFF | | | 19 | | 2 min | | 0 | 30 | 30 | 20 | 30 | No |  |
| Ferri ON | | | 20 | | 2 min | | 100 | 30 | 30 | 20 | 30 | No |  |
| water Rinse | | | 21 | | 40 sec | | 250 | 0 | 0 | 0 | 30 | yes |  |
| water Rinse | | | 22 | | 40 sec | | 250 | 0 | 0 | 0 | 30 | yes |  |
| water Rinse | | | 23 | | 40 sec | | 250 | 0 | 0 | 0 | 30 | yes |  |
| TCH ON | | | 24 | | 2 min | | 100 | 30 | 30 | 20 | 30 | yes |  |
| TCH OFF | | | 25 | | 2 min | | 0 | 30 | 30 | 20 | 30 | No |  |
| TCH ON | | | 26 | | 2 min | | 100 | 30 | 30 | 20 | 30 | No |  |
| TCH OFF | | | 27 | | 2 min | | 0 | 30 | 30 | 20 | 30 | No |  |
| TCH ON | | | 28 | | 2 min | | 100 | 0 | 0 | 20 | 30 | No |  |
| TCH OFF | | | 29 | | 2 min | | 0 | 30 | 30 | 20 | 30 | No |  |
| TCH ON | | | 30 | | 2 min | | 100 | 30 | 30 | 20 | 30 | No |  |
| water Rinse | | | 31 | | 0 | | 250 | 0 | 0 | 0 | 30 | yes |  |
| water Rinse | | | 32 | | 0 | | 250 | 0 | 0 | 0 | 30 | yes |  |
| water Rinse | | | 33 | | 0 | | 250 | 0 | 0 | 0 | 30 | yes |  |
| OsO4 2 ON | | | 34 | | 2 min | | 100 | 30 | 30 | 20 | 30 | yes |  |
| OsO4 2 OFF | | | 35 | | 2 min | | 0 | 30 | 30 | 20 | 30 | No |  |
| OsO4 2 ON | | | 36 | | 2 min | | 100 | 30 | 30 | 20 | 30 | No |  |
| OsO4 2 OFF | | | 36 | | 2 min | | 0 | 30 | 30 | 20 | 30 | No |  |
| OsO4 2 ON | | | 37 | | 2 min | | 100 | 30 | 30 | 20 | 30 | No |  |
| OsO4 2 OFF | | | 38 | | 2 min | | 0 | 30 | 30 | 20 | 30 | No |  |
| OsO4 2 ON | | | 39 | | 2 min | | 100 | 30 | 30 | 20 | 30 | No |  |
| water Rinse | | | 40 | | 40 sec | | 250 | 0 | 0 | 0 | 30 | yes |  |
| water Rinse | | | 41 | | 40 sec | | 250 | 0 | 0 | 0 | 30 | yes |  |
| water Rinse | | | 42 | | 40 sec | | 250 | 0 | 0 | 0 | 30 | yes |  |
| **Overnight in 2% AcU 4°C then heat 10 min at 37°C (without washing)** | | | | | | | | | | | | | 43 |

| **Day 2** | | | | | | | | |
| --- | --- | --- | --- | --- | --- | --- | --- | --- |
| **Step name** | **step number** | **time min** | **watts** | **vent time** | **vac time** | **set vac** | **load cooler T°** | yes |
| AcU 2 ON | 1 | 2 min | 100 | 30 | 30 | 20 | 30 | yes |
| AcU 2 OFF | 2 | 2 min | 0 | 30 | 30 | 20 | 30 | No |
| AcU 2 ON | 3 | 2 min | 100 | 30 | 30 | 20 | 30 | No |
| AcU 2 OFF | 4 | 2 min | 0 | 30 | 30 | 20 | 30 | No |
| AcU 2 ON | 5 | 2 min | 100 | 30 | 30 | 20 | 30 | No |
| AcU 2 OFF | 6 | 2 min | 0 | 30 | 30 | 20 | 30 | No |
| AcU 2 ON | 7 | 2 min | 100 | 30 | 30 | 20 | 30 | No |
| water Rinse | 8 | 40 sec | 250 | 0 | 0 | 0 | 30 | yes |
| water Rinse | 8 | 40 sec | 250 | 0 | 0 | 0 | 30 | yes |
| water Rinse | 9 | 40 sec | 250 | 0 | 0 | 0 | 30 | yes |
| Lead aspartate ON | 10 | 2 min | 100 | 30 | 30 | 20 | 30 | yes |
| Lead aspartate OFF | 11 | 2 min | 0 | 30 | 30 | 20 | 30 | No |
| Lead aspartate ON | 12 | 2 min | 100 | 30 | 30 | 20 | 30 | No |
| Lead aspartate OFF | 13 | 2 min | 0 | 30 | 30 | 20 | 30 | No |
| Lead aspartate ON | 14 | 2 min | 100 | 30 | 30 | 20 | 30 | No |
| Lead aspartate OFF | 15 | 2 min | 0 | 30 | 30 | 20 | 30 | No |
| Lead aspartate ON | 16 | 2 min | 100 | 30 | 30 | 20 | 30 | No |
| water Rinse | 17 | 40 sec | 250 | 0 | 0 | 0 | 30 | yes |
| water Rinse | 18 | 40 sec | 250 | 0 | 0 | 0 | 30 | yes |
| water Rinse | 19 |  |  |  |  |  |  |  |
| Acetonitrile 50% | 20 | 40 sec | 250 | 0 | 0 | 0 | 30 | yes |
| Acetonitrile 70% | 21 | 40 sec | 250 | 0 | 0 | 0 | 30 | yes |
| Acetonitrile 80% | 22 | 40 sec | 250 | 0 | 0 | 0 | 30 | yes |
| Acetonitrile 90% | 23 | 40 sec | 250 | 0 | 0 | 0 | 30 | yes |
| Acetonitrile 100% | 24 | 40 sec | 250 | 0 | 0 | 0 | 30 | yes |
| Acetonitrile 100% | 25 | 40 sec | 250 | 0 | 0 | 0 | 30 | yes |
| Acetonitrile 100% | 26 | 40 sec | 250 | 30 | 30 | 20 | 30 | yes |
| Resin 50 | 27 | 3 min | 150 | 30 | 30 | 20 | 30 | yes |
| Resin 75 | 28 | 3 min | 150 | 30 | 30 | 20 | 30 | yes |
| Resin 100 | 29 | 3 min | 150 | 30 | 30 | 20 | 30 | yes |
| Resin 100 | 30 | 3 min | 150 | 30 | 30 | 20 | 30 | yes |
| Resin 100 | 31 | 3 min | 150 | 30 | 30 | 20 | 30 | yes |
